# Single-cell multimodal profiling of pan-cancer cell lines uncovers gene regulatory principles underlying intrinsic cell states and environmental features

**DOI:** 10.64898/2026.05.31.729161

**Authors:** Zihan Xu, Aileen Ugurbil, Joshua Kwan, Chloe Schaefer, Abdulraouf Abdulraouf, Ziyu Lu, Erting Tang, Wei Zhou, Junyue Cao

## Abstract

Cancer arises from extensive genetic and epigenetic alterations that reshape chromatin, transcriptional regulation, and malignant cell states. To systematically chart cancer-intrinsic regulatory programs, we constructed a pan-cancer single-cell transcriptomic and epigenomic atlas encompassing 60 human cell lines representing 16 tissue origins and 20 cancer types, comprising 240,957 single-nucleus RNA-seq and 223,347 single-nucleus ATAC-seq profiles. Integrative analyses revealed extensive pan-cancer cell-state heterogeneity, core gene-regulatory networks, and a conserved epithelial-mesenchymal transition (EMT) axis that transcends tissue of origin. Copy-number variation analysis identified transcription factor amplification and downstream hyperactivation as key drivers of cancer cell-state reprogramming. To further examine how regulatory programs diverge within a cancer lineage and contribute to clinically divergent outcomes, we performed a focused comparison of cutaneous melanoma with acral melanoma, a rare, UV-independent subtype underrepresented in existing pan-cancer atlases. The comparison uncovered a universal inflammation-suppressive program in acral melanoma and an inflamed regulatory landscape in cutaneous melanoma, with the JAK-STAT pathway and downstream transcriptional responses as central discriminators. Integration of single-cell and bulk datasets across models and patient cohorts further linked in vitro tumor-intrinsic gene regulations with in vivo microenvironmental composition and immunotherapy responses. Together, by extending single-cell multi-omic profiling to rare alongside common cancer subtypes, this atlas offers a resource for mapping pan-cancer and subtype-specific gene-regulatory programs that shape cancer cell-state plasticity.

## Main

Gene regulation shapes cellular behavior, state transitions, and lineage development. Across chromatin and RNA layers, transcription factors (TFs) bind enhancers and promoters to modulate chromatin state and tune downstream gene expression^1^. Reconstructing gene regulatory networks (GRNs), which capture the interconnected regulatory architecture through which TFs coordinate gene-expression programs, can therefore provide mechanistic insight into the regulation of cellular phenotypes^2,3^. Cancer cell states are shaped by extensive epigenetic and transcriptional remodeling that drives their aberrant behaviors^4^, yet the core regulatory programs shared across diverse malignancies remain poorly defined. Large-scale cancer genomics studies have cataloged molecular features such as DNA mutations and bulk expression profiles across cancer types^5,6^, but bulk profiling masks regulatory heterogeneity among malignant cells and between malignant and non-malignant cells within tumors. Although recent single-cell studies have begun to resolve cellular heterogeneity in cancer^7–9^, many have focused on individual cancer types, single modalities such as gene expression, or relatively limited sample cohorts, restricting the ability to reconstruct regulatory circuits across epigenetic and transcriptional layers.

Cancer cell lines provide a uniquely tractable system in which tumor-intrinsic programs can be isolated from microenvironmental influences, enabling precise dissection of core gene regulatory networks across heterogeneous cancer lineages. While recent single-cell atlases of cancer cell lines have revealed extensive molecular heterogeneity^10–12^, most studies rely on droplet-based platforms that are restricted to a single molecular modality or limited cell numbers per experiment, constraining the ability to map chromatin–gene regulatory relationships across diverse cancer states. Overcoming these limitations requires a large-scale, multimodal analysis of heterogeneous cancer cell lineages, which would enable the identification of lineage-specific and convergent regulatory programs, an essential step toward defining shared intrinsic regulatory logic across malignancies.

To address this gap, we used EasySci^13^, a combinatorial-indexing single-cell genomics platform that offers high scalability and sample-multiplexing capability, to construct a pan-cancer single-cell atlas comprising 60 human cancer cell lines representing 16 tissue origins and 20 cancer types. This dataset expands prior cancer cell line atlases by an order of magnitude, profiling more than 460,000 transcriptomic and chromatin-accessibility profiles. Through integrative analysis of these multimodal data, we uncovered a pan-cancer epithelial–mesenchymal transition (EMT) axis linked to transcription factor amplification, as well as a striking regulatory divergence between acral and cutaneous melanoma that maps to distinct immunological features and immunotherapy response in patients. These findings demonstrate that cancer-intrinsic regulatory logic captured in vitro can be predictive of clinical outcomes, thereby establishing a framework for linking single-cell regulatory programs to precision oncology.

## Results

### Single-cell multi-omic atlas of diverse human cancer cell lines

To uncover cell-state heterogeneity and gene regulatory networks underlying distinct cancer cell states, we applied EasySci^13^ for single-cell RNA-seq and ATAC-seq analysis across sixty human cancer cell lines spanning twenty cancer types, including skin cutaneous melanoma, acral melanoma, breast cancer, colon adenocarcinoma, lung cancer, kidney renal cell carcinoma, hepatocellular carcinoma, and tumors associated with the hematopoietic and lymphatic systems, among others (**Figure 1a-b**). Based on combinatorial indexing, EasySci enables parallel sample barcoding through indexed reverse transcription of different samples in separate wells of a plate. Subsequent split-pool barcoding exponentially expands the cell barcode space, enabling profiling of millions of cells in a single experiment. These features simplify multi-sample profiling and minimize technical batch effects^14^. Compared with previous single cell multi-omic analysis of cancer cell atlas that mostly focused on breast cancer and colorectal cancer^10^, our panel includes multiple cell lines for acral melanoma (WM3211, WM4235, WM4324, YUHIMO, M040204, M040416, MB4667, YUSEEP, YUSUSA, M141207, M160113), cutaneous melanoma (A375, SK-Mel-24, SK-Mel-3, MeWo, RPMI7951), and lung adenocarcinoma (H2030-TGL, H2030-BrM3, PC9-TGL, PC9-BrM3, H2087-TGL, H2087-LCC1, H2087-LCC2), thus enabling the identification of gene regulatory circuits shared among cells of the same tumor types.

**Fig. 1.**
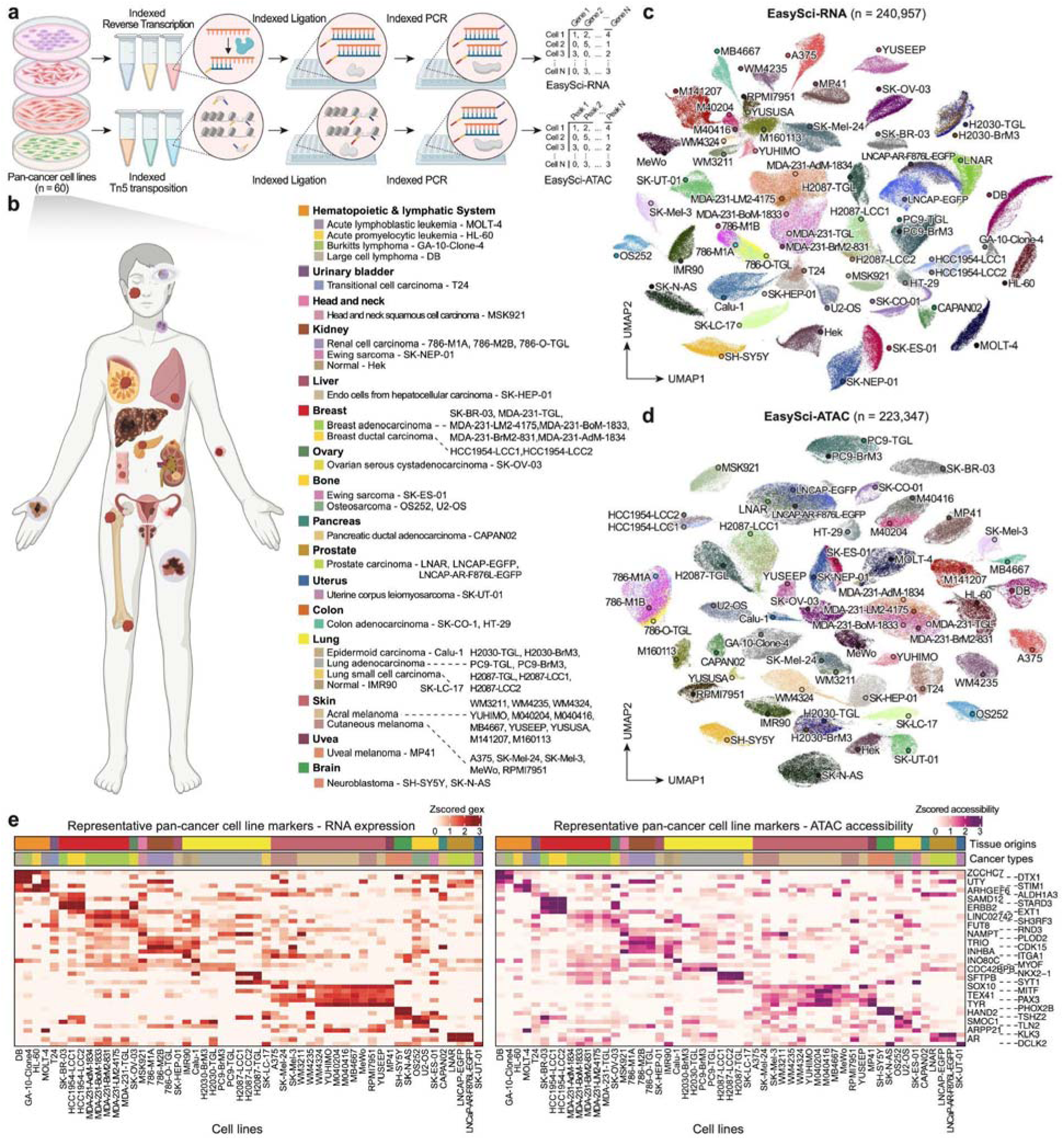
EasySci enables parallel transcriptome and chromatin accessibility profiling across diverse cancer cell lines. **a,** Schematic of the EasySci-RNA and EasySci-ATAC library preparation workflows. Cancer cell lines were profiled separately for transcriptome and chromatin accessibility using combinatorial indexing strategies. **b,** Overview of the 60 profiled cell lines, representing 20 cancer types and two non-malignant cell lines, a fetal lung fibroblast line (IMR-90) and an embryonic kidney epithelial cell line (HEK293). Notably, the panel included 11 acral melanoma cell lines. c-d, UMAP visualization of 240,957 single cells profiled by EasySci-RNA (c) and 223,347 single cells profiled by EasySci-ATAC (d), colored by cell line identity, with each cluster annotated by its corresponding cell line name. Cells cluster primarily by cancer type and tissue of origin. **e,** Representative marker genes show concordant RNA expression and ATAC gene-body accessibility patterns specific to each cancer’s tissue of origin.

Nuclei were isolated from each cancer cell line and distributed into indexed wells for reverse transcription (EasySci-RNA) or transposition (EasySci-ATAC), allowing the cell identity to be directly retrieved based on each cell’s barcode. Following library preparation and sequencing, we recovered 240,957 single-nucleus transcriptomes (median 3,784 nuclei per cell line; median 2,351 unique transcripts per cell; **Extended Data Fig. 1a, b, e**) and 223,347 single-nucleus chromatin-accessibility profiles (median 3,768 nuclei per cell line; median 7,192 unique fragments per cell; **Extended Data Fig. 1c, d, f**).

**Extended Data Fig. 1.**
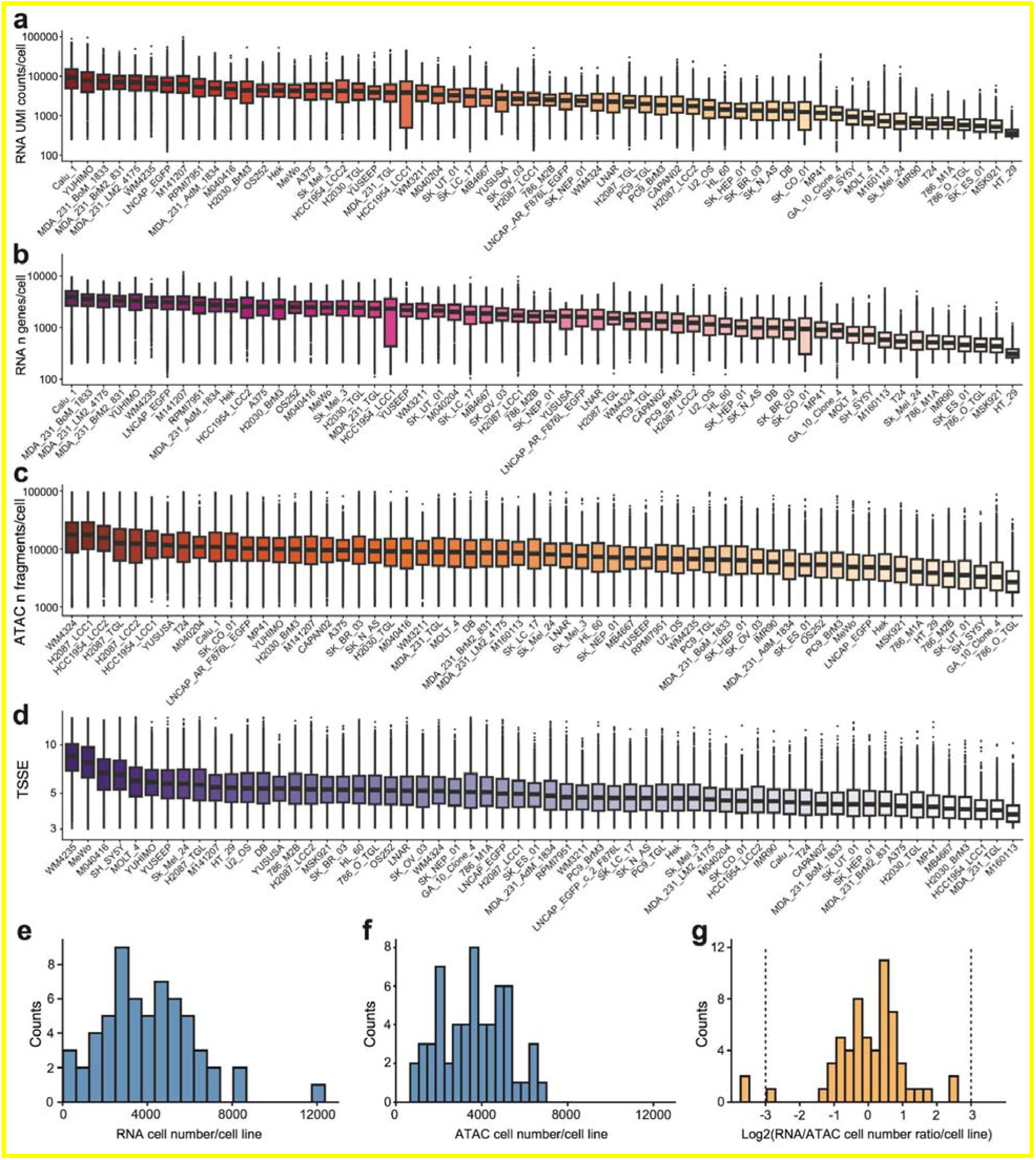
Quality metrics of pan-cancer cell line multi-modal single-cell profiling. **a,** Box plot showing the distribution of single-cell RNA UMI counts across cell lines. **b,** Box plot showing the distribution of the number of genes detected per single cell across cell lines. **c,** Box plot showing the distribution of the number of ATAC fragments per single cell across cell lines. **d,** Box plot showing the distribution of single-cell TSSE scores across cell lines. **e,** Histogram showing the distribution of single-cell RNA-seq cell numbers across cell lines. **f,** Histogram showing the distribution of single-cell ATAC-seq cell numbers across cell lines. **g,** Histogram showing the distribution of log2 RNA-to-ATAC cell numbers across cell lines. Dashed lines indicate the cutoffs used for filtering. Only cell lines without extreme imbalance between RNA and ATAC cell numbers were retained for integration.

UMAP embeddings of RNA or ATAC profiles showed that cells clustered primarily by cancer type and tissue of origin (**Figure 1c-d**). In addition, representative marker genes of tissues exhibited concordant RNA expression and gene-body accessibility activations in subsets of cancer cell lines, such as *TYR* and *PAX3* in skin melanoma cells, which support melanogenesis and melanocytic/neural crest lineage identity^15^; *NKX2-1* and *SFTPB* in lung carcinoma cells, which define lung epithelial identity and surfactant production^16,17^; *STARD3* and *ERBB2* in HER2-positive breast cancer cells, which mark the HER2-amplified locus and HER2 signaling^18^; *KLK3* and *AR* in prostate cancer cells, which encode prostate-specific antigen and the androgen receptor transcription factor, respectively^19^; and *PHOX2B* and *HAND2* in neuroblastoma cells, which define sympathetic neuronal lineage identity^20^ (**Figure 1e**). These results indicate that, despite being cultured under similar in vitro conditions, the cancer cell lines retain the gene expression and epigenetic signatures of their tissues of origin.

### Pan-cancer GRN analysis defines EMT-linked cell states shared across lineages

To dissect the transcriptional regulatory features across these phenotypically diverse cancer cells, we sought to identify cis-regulatory elements (CREs) of genes and potential upstream TF regulators (**Methods**). EasySci-RNA and EasySci-ATAC profile transcriptomes and chromatin accessibility in separate cells, so no single cell carries both measurements. Given the balanced coverage across cell lines and RNA/ATAC modalities (**Extended Data Fig. 1g**), we performed computational integration across modalities using a meta-cell strategy. In this approach, single-cell transcriptomes and chromatin accessibility profiles were first integrated into a shared low-dimensional space, after which transcriptionally and epigenetically similar neighboring cells were identified and grouped to form ‘meta-cells’. Each meta-cell therefore represents the multi-modal profile of a local neighborhood of cells. By aggregating counts from locally similar cells, this approach links transcriptomic and chromatin states, reduces dropout-driven sparsity at each measurement point, and preserves cell-state heterogeneity at the meta-cell level (**Fig. 2a, Extended Data Fig. 2**). We then used these multi-modal meta-cells to identify ATAC peak–gene expression associations, perform TF motif scanning, and define regulons, here referring to groups of target genes co-regulated by a TF, to construct a pan-cancer GRN (**Fig. 2a**). Following rigorous data curation, consistent with prior studies^10,12^, we observed that cell-cycle status dominated intra-cell line heterogeneity at the multimodal level in most cases, whereas more discrete cell states were also robustly detected in specific cell lines (**Extended Data Fig. 2a-j**). With pan-cancer GRN construction, altogether, we identified 85,670 linkages between 61,259 ATAC peaks and 13,295 target genes, with a median of 5 CREs regulating a gene (**Extended Data Fig. 3a-c**). Downsampling analyses further supported the importance of sufficient cell coverage and the robustness of the network construction (**Extended Data Fig. 3d-f**).

**Fig. 2.**
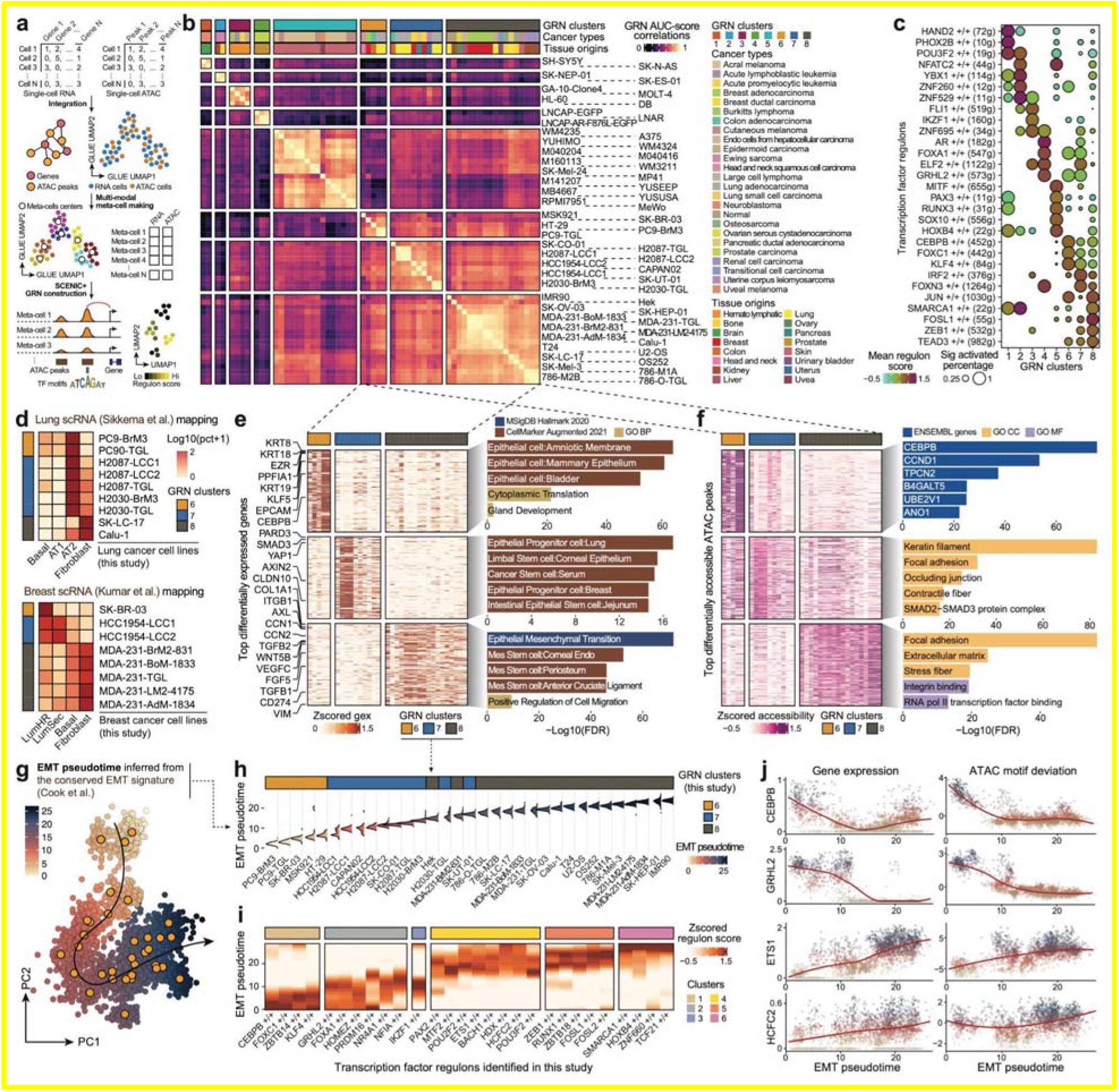
Pan-cancer gene regulatory networks delineate intra- and cross-lineage molecular states *de novo*. **a,** Schematic of pan-cancer gene-regulatory network construction. Single-cell RNA-seq and ATAC-seq data were integrated, then neighboring cells were aggregated into multimodal meta-cells. GRNs were built by associating TFs with enhancers and linking enhancers to nearby target genes. Activity scores of the resulting regulons were then computed across meta-cells. **b,** Heatmap, ordered by hierarchical clustering, showing correlations of TF-regulon AUCell scores across cell lines (n=59). Annotations indicate GRN clusters, cancer types, and tissue origins. **c,** Dot heatmap showing representative TF regulons activated in cell lines across GRN clusters. Dot size denotes the proportion of cell lines within a cluster with significant regulon activation; color denotes the mean gene-program expression score across cell lines in the cluster. **d,** Heatmaps showing the proportion of cells from lung cancer (top) and breast cancer (bottom) mapped to phenotypically similar normal cell types from references. **e,** Heatmap showing the top differentially expressed genes across GRN cluster 6-8 with highly mixed cancer types and tissue origins; the right panel shows the enrichment of functional terms for each group. **f,** Heatmap showing the top differentially accessible ATAC peaks across GRN cluster 6-8 with highly mixed cancer types and tissue origins; the right panel shows the enrichment of functional terms for genes nearest the peaks in each group. **g,** PCA plot showing meta-cell distribution along the EMT. Meta-cells are colored by inferred pseudotime; orange points (n = 33) mark the meta-cell cluster centers for each cell line in PCA space. The trend line is fitted using slingshot^40^. **h,** Ridge plot showing the distribution of EMT pseudotime across cell lines within GRN cluster 6-8. The top annotation bar shows the orthogonal-clustering identity of the cell lines. **i,** Heatmap with hierarchical clustering of gene program scores for EMT-associated TF regulons across the trajectory. **j,** Scatter plots with fitted trend lines showing coordinated gene expression and TF motif deviation across the EMT pseudotime of meta-cells. Points are colored by assigned pseudotime. Panel d uses external reference single-cell atlases of normal lung and breast tissue. The EMT trajectory in panels g-j is defined by a published conserved EMT signature applied to our cells. Panel a is a schematic; all other panels show analyses generated in this study.

**Extended Data Fig. 2.**
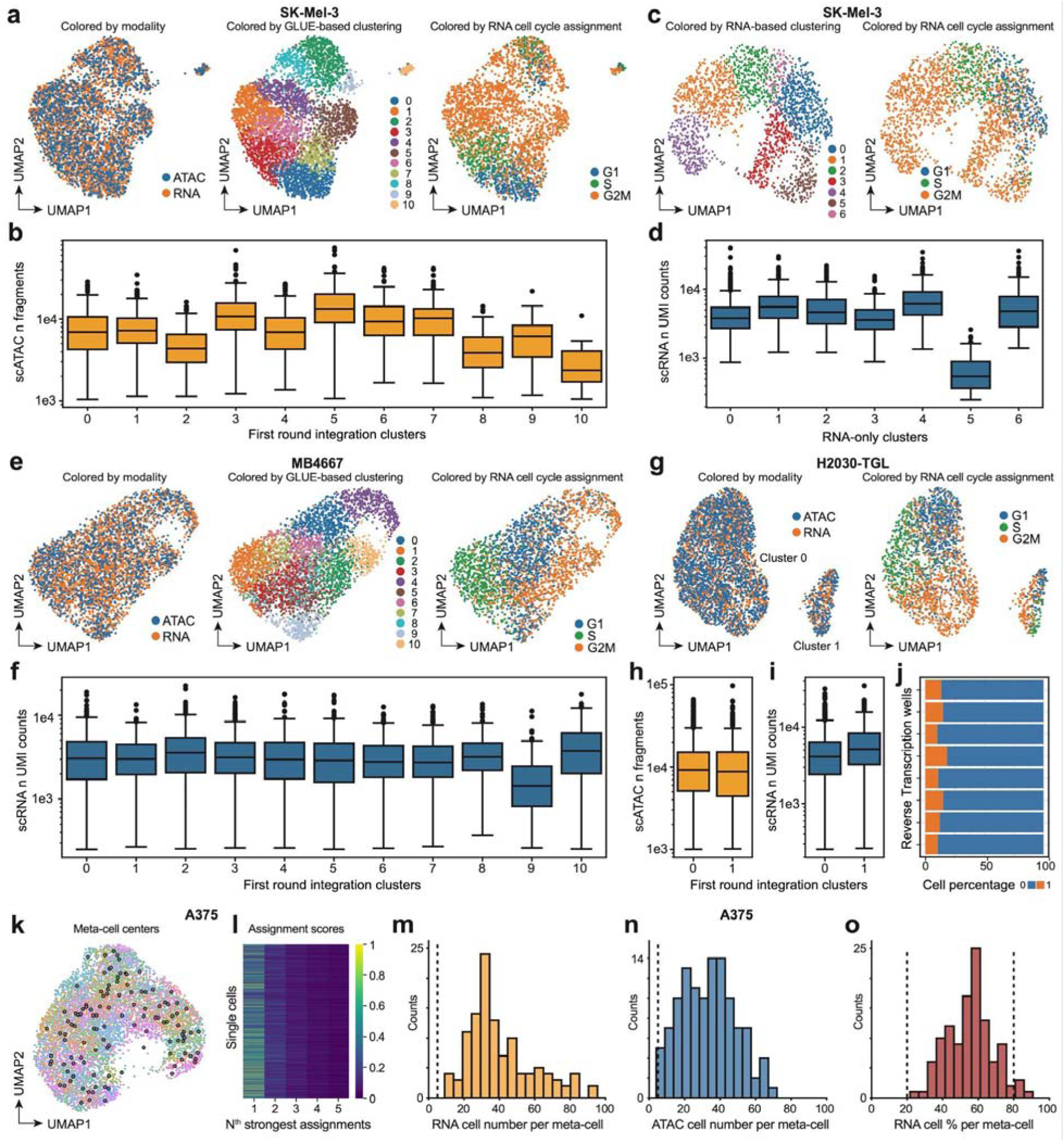
Integration and processing of single-cell RNA and ATAC and meta-cell identification. **a,** Example of low-quality cell filtering driven by the ATAC modality: first-round GLUE integration-based UMAPs of SK-Mel-3 cells colored by RNA/ATAC modalities (left), Leiden clusters (middle), and RNA-based cell cycle phase assignments (right). **b,** Box plot showing the distributions of single-cell fragment numbers across Leiden clusters of SK-Mel-3. Outlier clusters (2, 8, 9, 10) with mixed cell cycle assignments were composed of low-quality ATAC cells. **c,** RNA gene expression–based UMAP of SK-Mel-3 cells colored by Leiden clusters (left) and cell cycle phase assignments (right), after removing low-quality cells and outlier clusters identified from first-round integration. **d,** Box plot showing the distributions of single-cell RNA UMI counts across gene expression–based Leiden clusters of SK-Mel-3. The outlier cluster (5) consisted of low-quality RNA cells. **e,** Example of low-quality cell filtering driven by the RNA modality: first-round GLUE integration-based UMAPs of MB4667 cells colored by RNA/ATAC modalities (left), Leiden clusters (middle), and RNA-based cell cycle phase assignments (right). **f,** Box plot showing the distributions of single-cell RNA UMI counts across Leiden clusters of MB4667. The outlier cluster (9) with mixed cell cycle assignments consisted of low-quality RNA cells. **g,** Example of a true discrete intra–cell line state: first-round GLUE integration-based UMAPs of H2030-TGL cells colored by RNA/ATAC modalities (left) and RNA-based cell cycle phase assignments (right). This discrete state in H2030-TGL was also observed in an independent study^41^. **h,** Box plot showing the distributions of single-cell ATAC fragment numbers across main and outlier clusters. The outlier could not be explained by low ATAC quality. **i,** Box plot showing the distributions of single-cell RNA UMI counts across main and outlier clusters. The outlier could not be explained by low RNA quality. **j,** Bar plot showing the RT-well composition of cells in both clusters. The outlier cluster could not be explained by batch effects during library preparation. **k,** Example of meta-cell identification: final GLUE integration-based UMAP of A375 cells colored by meta-cell assignments. The centers of meta-cells are highlighted with black-circled dots. **l,** Heatmap showing the top five SEACells assignment scores for all A375 single cells. **m,** Histogram showing the distribution of the number of RNA cells per meta-cell identified from A375 integration. **n,** Histogram showing the distribution of the number of ATAC cells per meta-cell identified from A375 integration. **o,** Histogram showing the distribution of the percentage of RNA cells within meta-cells identified from A375 integration. Boxes in box plots indicate the median and interquartile range (IQR), with whiskers indicating 1.5× IQR.

**Extended Data Fig. 3.**
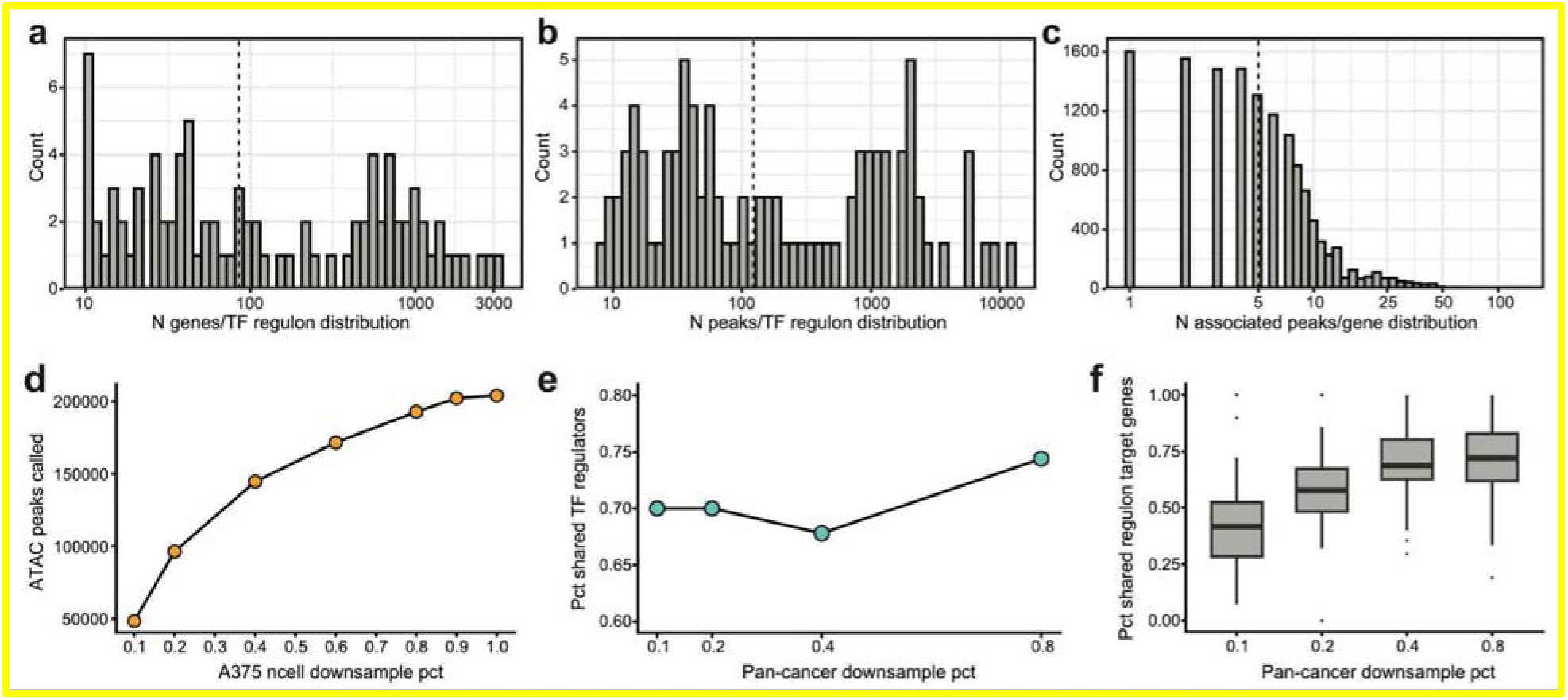
Evaluation of the robustness of pan-cancer gene regulatory network construction. **a,** Histogram showing the distribution of the number of target genes per TF. The dashed line indicates the median (85). **b,** Histogram showing the distribution of the number of ATAC peaks regulated by TFs. The dashed line indicates the median (122.5). **c,** Histogram showing the distribution of the number of ATAC peaks associated with downstream genes. The dashed line indicates the median (5). **d,** Number of ATAC peaks called from the A375 cell line across increasing cell downsampling fractions. **e,** Fraction of TF regulators identified to be shared with the full dataset across pan-cancer downsampling fractions. **f,** Fraction of regulon target genes identified to be shared with the full dataset across pan-cancer downsampling fractions. Boxes in box plots indicate the median and interquartile range (IQR), with whiskers indicating 1.5× IQR.

To classify cell lines by their regulatory programs, we focused on activating regulons, defined as those in which TF expression is positively associated with target gene expression, and the accessibility of TF motif-containing cis-regulatory elements is also positively associated with target gene expression (**Extended Data Fig. 4a-b**). Scoring pan-cancer cell lines with these 92 activating TF regulons, followed by hierarchical clustering, identified 8 clusters of cancer cell lines with distinct gene regulatory programs (**Fig. 2b**), in which 5 (Cluster 1-5) were defined by the activity of tissues- and lineage-specific TFs. Cluster 1, consisting of SH-SY5Y and SK-N-AS neuroblastoma cell lines, exhibited strong activation of PHOX2B and HAND2 regulation, both well-established core factors in neurogenesis and neuroblastoma^20^. Cluster 2, consisting of the Ewing sarcoma cell lines SK-ES-01 and SK-NEP-01, showed high activity of YBX1, which has been reported to be essential for the malignant behavior of sarcoma^21^. Cluster 3 contained four hematopoietic and lymphatic system malignancy cell lines, including three lymphocytic cell lines (GA-10-clone4, MOLT-4, DB) and one promyelocytic leukemia cell line (HL-60), and was characterized by IKZF1, an essential transcription factor in lymphocytic differentiation^22^. Notably, its aberrant activation in HL-60 reflected impaired myeloid differentiation^23^. Cluster 4, composed of prostate cancer cell lines, exhibited strong activation of AR and FOXA1, the master regulators of androgen signaling and prostate lineage identity^19^, while cluster 5, composed of melanoma cell lines, showed high activity of key melanocytic lineage-determining transcription factors, including MITF, SOX10, and PAX3^15^.

**Extended Data Fig. 4.**
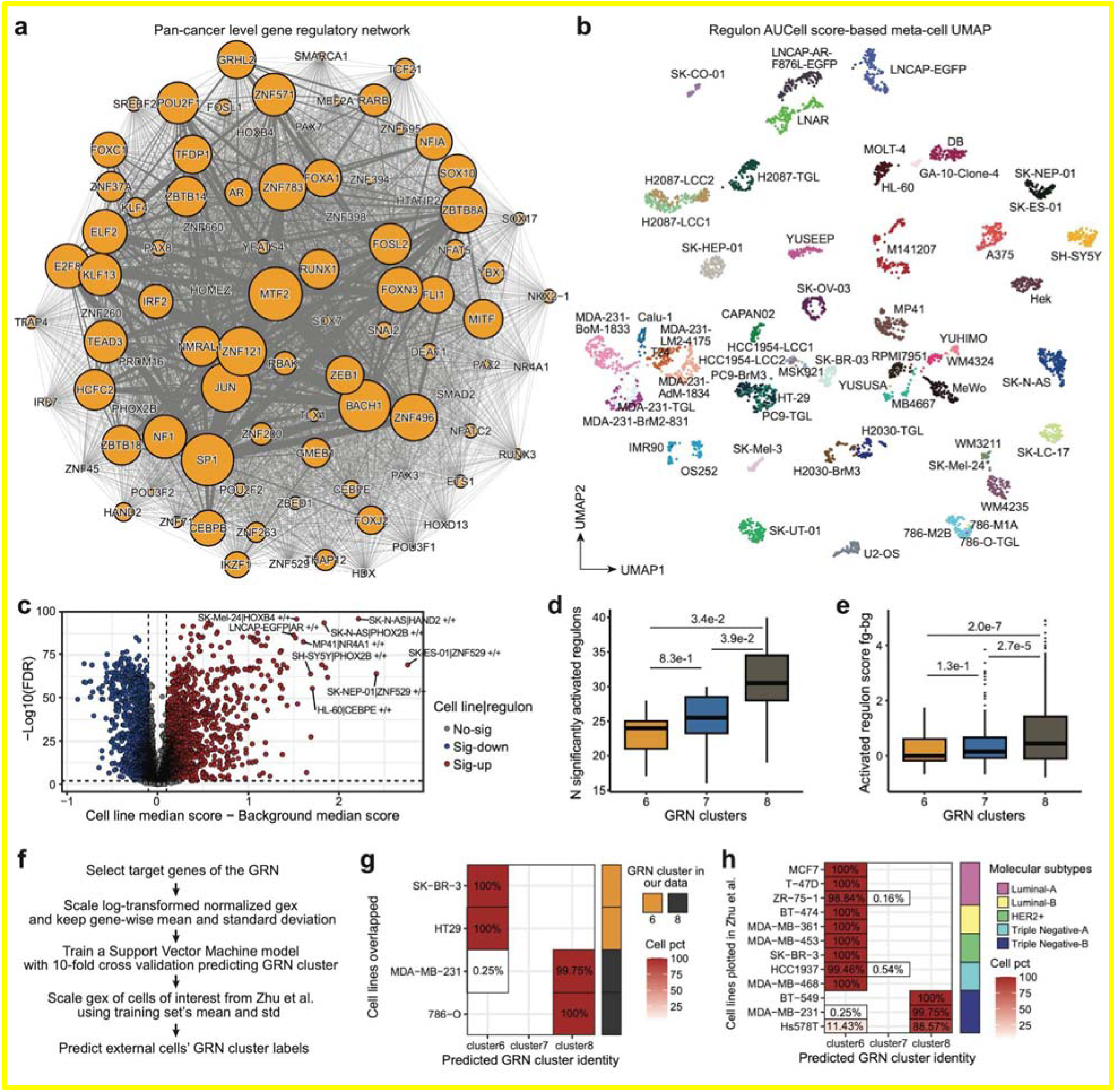
Characteristics of the Pan-cancer gene regulatory network. **a,** Network diagram showing pan-cancer TF regulons. Dot size represents the number of target genes for each TF, and edge width represents the number of shared target genes between two regulons. **b,** UMAP of pan-cancer cell line meta-cells constructed from regulon activity scores (AUCell) for nearest-neighbor graph reconstruction. **c,** Volcano plot showing the relative activation and statistical significance of TF regulons across cancer cell lines. The most strongly activated regulons exhibited high tissue specificity, such as HAND2 and PHOX2B in neuroblastoma, AR in prostate cancer, HOXB4 in melanoma, and CEBPE in acute promyelocytic leukemia. **d,** Box plot showing the distributions of the number of significantly activated TF regulons across cancer cell lines within GRN clusters 4-6, corresponding to epithelial, intermediate, and mesenchymal states. Multiple test-corrected p-values from Tukey test following one-way ANOVA were reported. **e,** Box plot showing the distributions of the extent of significantly activated TF regulons across cancer cell lines within GRN clusters 4-6, corresponding to epithelial, intermediate, and mesenchymal states. The difference between the z-scored regulon score of a cancer cell line and the mean background z-scored regulon score was used as a proxy. Multiple test-corrected p-values from Tukey test following one-way ANOVA were reported. **f,** A brief workflow describing the establishment of a validation model for predicting GRN-based cancer cell line phenotypes. **g,** Predicted GRN cluster identities of four overlapping cell lines between our dataset and Zhu et al., showing assignment to the expected epithelial cluster 6 or mesenchymal cluster 8. **h,** Predicted GRN cluster identities of breast cancer cell lines from Zhu et al., showing concordance between epithelial/stromal-EMT phenotypes and assignment to GRN cluster 6 or 8. Boxes in box plots indicate the median and interquartile range (IQR), with whiskers indicating 1.5× IQR.

In contrast, clusters 6–8 contained a diverse mixture of cancer cell lines from distinct tissues and cancer types (**Fig. 2b**), including those from breast, lung, skin, head and neck, kidney, bone, colon, ovary, and uterus. This strongly suggests the existence of universal cell states that share similar gene-regulatory features beyond lineage specificity. The activation of TF regulons displayed a continuous transition across these clusters (**Fig. 2c**). For example, the activity of TFs that peaked in cluster 6, such as CEBPB and GRHL2 declined in clusters 7 and 8, while most TF regulons highly active in cluster 8, including FOSL1, ZEB1, TEAD3 that has been characterized to be involved in cancer dedifferentiation^24^ and EMT^25^, were relatively inactive in clusters 6 and 7. Furthermore, compared with clusters 6 and 7, cell lines in cluster 8 exhibited both the largest number of active TF regulons and the strongest overall transcriptional activation (**Extended Data Fig. 4c-e**), recapitulating the hypertranscriptional property of aggressive cancers^26,27^ and the stemness-associated transcriptional state^28^.

To further dissect the phenotypes of these pan-cancer conserved cell states, we implemented a nearest-neighbor-based annotation label transfer strategy. In brief, query cells from lung and breast cancer cell lines within clusters 6–8 were projected into a shared transcriptional space with reference single-cell RNA-seq atlases of normal lung^29^ and breast tissues^30^. Cell-type annotations were then assigned to query cancer cells based on the labels of their nearest neighbors among the reference cells (**Fig. 2d, Methods**). Consistent with the developmental scheme in which alveolar type 2 (AT2) cells serve as progenitors that differentiate into terminal alveolar type 1 (AT1) cells, and with recent findings^31^ identifying AT2 as the origin of lung adenocarcinoma, most lung cancer cell lines in clusters 6 and 7 originated from AT2 cells. In contrast, cluster 8 cells were mapped almost completely to a fibroblast phenotype (**Fig. 2d, top**). In the mammary tissue, luminal hormone-responsive (LumHR) cells represent a differentiated hormone-sensing state, whereas luminal secretary (LumSec) cells retain a more progenitor-like identity. A similar phenotypic transition was observed in breast cancer cell line mapping: SK-BR-3 mapped mainly to LumHR, consistent with a recent finding identifying LumHR cells as the origin of ER⁺ breast cancer^32^, while HCC1954 mapped to both LumHR and LumSec, reflecting higher plasticity. In contrast, triple-negative MDA-MB lines in cluster 8 exhibited a complete mesenchymal phenotype (**Fig. 2d, bottom**). Overall, the *in vitro* cancer cell lines from both tissues retained transcriptomic features from their cell type of origin, and strikingly, their phenotypic transition across clusters 6–8 recapitulated the progression from specialized epithelial cells to a fibroblast-like mesenchymal state.

At the pan-cancer level, analysis of cluster-specific activated genes and ATAC peaks further supported these phenotypic assignments (**Methods**). Functional enrichment of differentially expressed genes confirmed that clusters 6–8 correspond to epithelial, intermediate, and mesenchymal states along the EMT trajectory (**Fig. 2e**). Consistently, accessible CREs in cluster 8 were enriched near genes associated with mesenchymal functions, such as stress fiber formation for stronger migratory ability of mesenchymal cells^33^. In contrast, the epithelial state showed strong chromatin activation at regulatory regions near genes such as *CEBPB*. Notably, ATAC peaks significantly activated in cluster 7 showed partial activation in both clusters 6 and 8, underscoring its role as an intermediate state along the EMT transition trajectory (**Fig. 2f**).

To further confirm the generalizability and robustness of our epigenetically and transcriptionally defined EMT-like states, we utilized a recently defined conserved EMT signature derived from diverse cell lines treated with pro-EMT cytokines^34^. This signature captures genes consistently induced across multiple EMT-promoting conditions and was used here as an orthogonal validation (**Methods**). Cells in our study aligned along a continuous trajectory defined by the published conserved EMT signature derived from distinct cancer cell lines^34^ (**Fig. 2g**), and their pseudotime distributions closely paralleled the GRN-based clustering independently defined by our epigenetic and gene expression association analysis (**Fig. 2h**). Further associating TF regulon activity to EMT progression revealed both established and new EMT regulators (**Fig. 2j, Methods**). In addition to more characterized EMT inducers (*e.g*., ZEB1^25^ and FOSL1^35,36^), ETS1 and HCFC2 also showed coordinated activation at the chromatin and transcriptional levels, accompanied by increased expression of their targets across pan-cancer EMT. Conversely, GRHL2, CEBPB, and FOXC1 showed epithelial-state-enriched expression and activity, with regulon activities negatively associated with EMT progression. These findings are consistent with previous studies in specific cancer models implicating GRHL2 in EMT suppression in breast, lung, and ovarian cancer, as well as CEBPB and FOXC1 in breast cancer^37–39^, and our analysis extends these cancer-type-specific observations to a broader pan-cancer context (**Fig. 2j**).

To validate the robustness of the regulatory phenotypes identified in our dataset, we trained a support vector machine model using the expression of pan-cancer GRN target genes to predict single-cell phenotype labels (**Extended Data Fig. 4f**). When applied to an external cancer cell-line single-cell transcriptomic dataset generated by Zhu et al.^10^, this model successfully recovered pan-cancer EMT states in both overlapping cell lines (**Extended Data Fig. 4g**) and previously unseen cell lines annotated by Zhu et al. as occupying more differentiated epithelial or more mesenchymal states (**Extended Data Fig. 4h**), demonstrating that the gene-regulatory programs we define, not only the transcriptional states, are reproducible across platforms.

In summary, we defined core GRNs across diverse cancer cells and identified both lineage-specific and lineage-independent EMT states governed by conserved epigenetic and transcriptomic features.

### Copy number variation modulate the epithelial-mesenchymal cell state continuum

As copy number variation (CNV) has been extensively recognized as a key driver of cancer cell behavior and is strongly associated with tumor response to therapies and patient prognosis^42^, we asked whether it also contributes to the cross-lineage cell state alterations. We inferred genome-wide amplification and deletion patterns at the meta-cell level using InferCNV^43^ (**Fig. 3a, Methods**). As expected, no significant subclones were detected within individual cell lines, consistent with their continuous clonal evolution in culture. Cell lines with closer lineage proximity also displayed more similar CNV landscapes (**Extended Data Fig. 5**). In agreement with previous reports, the karyotypically normal diploid cell line IMR90 showed minimal CNVs (**Extended Data Fig. 5a**), and our inferCNV profiles were highly concordant with whole-exome- and whole-genome-sequencing-derived CNV patterns from independent studies^44–46^ (**Extended Data Fig. 5c-e**). By systematically examining the proportion of amplified genes among TFs with regulon activation and among member genes of these regulons across all cell lines, we found that TFs were amplified more frequently than their downstream targets (**Fig. 3b**), suggesting that TF amplification and hyperactivation could represent a common mechanism governing cancer cell state heterogeneity.

**Fig. 3.**
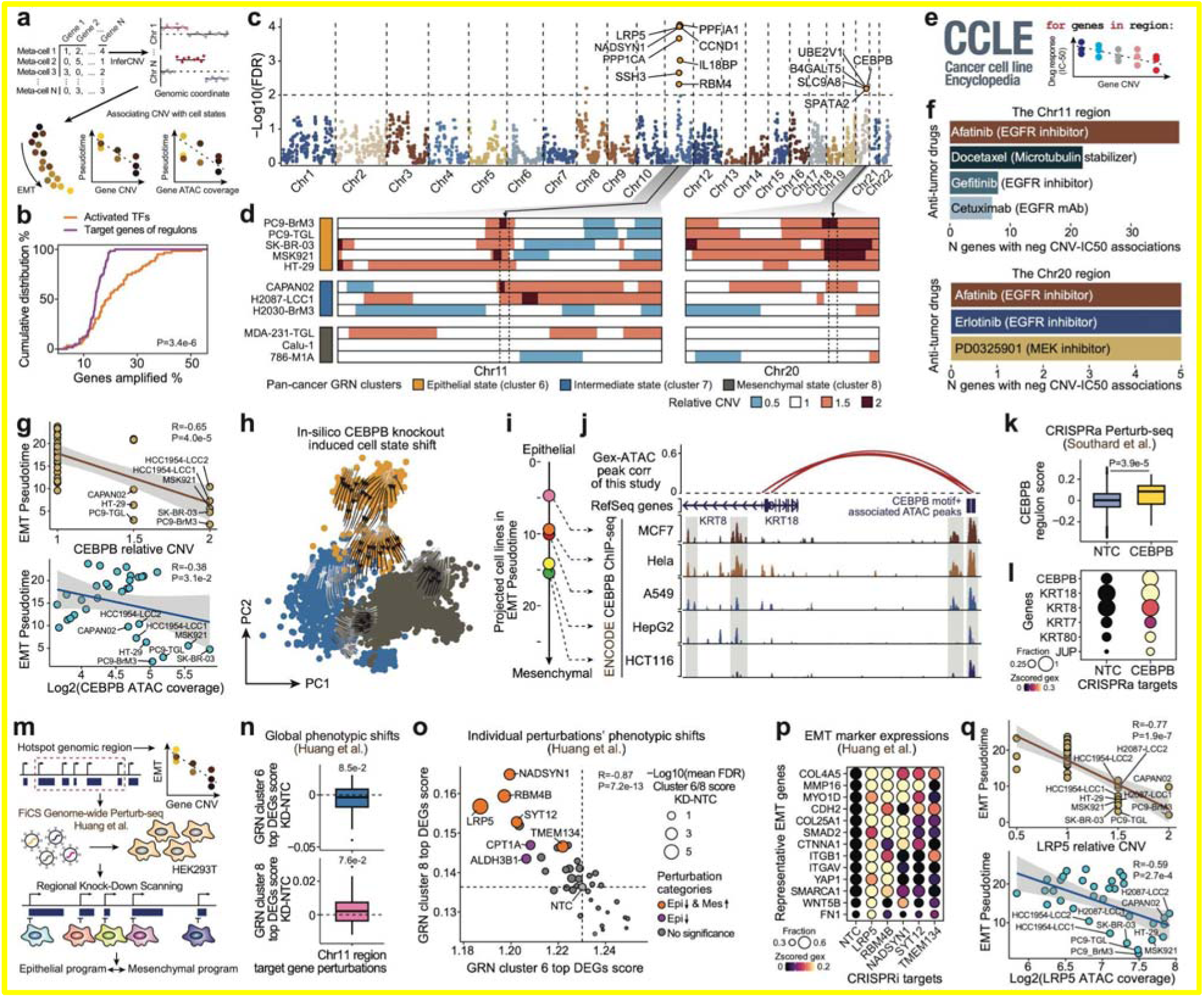
Systematic examination of pan-cancer copy number variation nominated key regulators of cell states. **a,** Schematic of pan-cancer copy-number–EMT association analysis and CNV calling. **b,** Cumulative distribution of the percentage of genes with amplifications in activated TF sets or among target genes of activated regulons across cancer cell lines. **c,** Scatter plot of associations between gene copy number and EMT state. Each dot is a gene ordered by genomic coordinates across the genome; only autosomes were considered. **d,** Heatmaps showing CNV patterns on chromosomes 11 and 20 for representative cell lines from GRN clusters 6–8, corresponding to epithelial, intermediate, and mesenchymal states. Dashed lines indicate regions with CNV patterns associated with EMT states. **e,** Schematic of the association analysis between CNV and drug sensitivity across CCLE cancer cell lines. **f,** Bar plots showing top antitumor drugs whose IC50 values across cell lines are negatively associated with genes within identified hotspot regions. **g,** Scatter plots with fitted lines showing negative associations between EMT pseudotime and CEBPB copy number (top) or normalized ATAC signal coverage (bottom). **h,** PCA plot with state-shift vectors indicating predicted future cell-state changes after in-silico CEBPB knockdown. Target-gene expression was modeled from TFs, and “future” transcriptomes were computed over ten iterations after setting CEBPB expression to 0. **i,** 1D dot plot showing median EMT pseudotime of external normal and cancer cell lines inferred by nearest-neighbor transfer. **j,** Representative genome tracks showing −log10 P values (CEBPB vs input) from ENCODE ChIP-seq across cell lines. Proximal ATAC peaks putatively regulated by CEBPB and associated with KRT8 and KRT18 are highlighted; peak–gene links are shown as arches. **k,** Box plot comparing single-cell gene-program scores of CEBPB regulons between NTC and CEBPB-activated cells. NTC cells n = 13,673, CEBPB CRISPRa cells n = 62. **l,** Dot heatmap showing z-scored expression of representative epithelial marker genes in NTC and CEBPB-activated cells that are also identified as CEBPB targets in the pan-cancer GRN. **m,** Schematic of examination of lead phenotype-associated genes within the chr11 hotspot region. **n,** Boxplots showing the global distribution of relative EMT state transitions compared to the Non-Targeting Control (NTC) upon knockdown of different genes within the chr11 region. The top 100 significantly upregulated genes of pan-cancer cell lines in GRN cluster 6 (epithelial state) and GRN cluster 8 (mesenchymal state) were used for phenotypic state scoring. P-values from one-sided one-sample Wilcoxon tests are reported. Number of targeted genetic perturbations: n = 39. **o,** Scatter plot showing the statistical significance and effect size of EMT state transitions for individual genetic perturbations using perturbed transcriptomes of HEK293T cells relative to the NTC. The gray dot represents the NTC. Multiple-test-corrected FDR values from two-sample Wilcoxon tests are reported. **p,** Dot heatmap showing relative expression of representative EMT marker genes under genetic perturbations that induced significant EMT state shifts, alongside the NTC. These genes were at least significantly upregulated in one of these perturbations compared to NTC. **q,** Scatter plots with fitted lines showing negative associations between EMT pseudotime and LRP5 copy number (top) or normalized ATAC signal coverage (bottom). Boxes in box plots indicate the median and interquartile range (IQR), with whiskers indicating 1.5× IQR. External datasets are used in panel f (CCLE drug sensitivity), i (external cell-line transcriptomes), j (ENCODE CEBPB ChIP-seq), k-l (CRISPRa Perturb-seq), and n-p (CRISPRi Perturb-seq). Panels a, e, and m are schematics; all other panels show analyses generated in this study.

**Extended Data Fig. 5.**
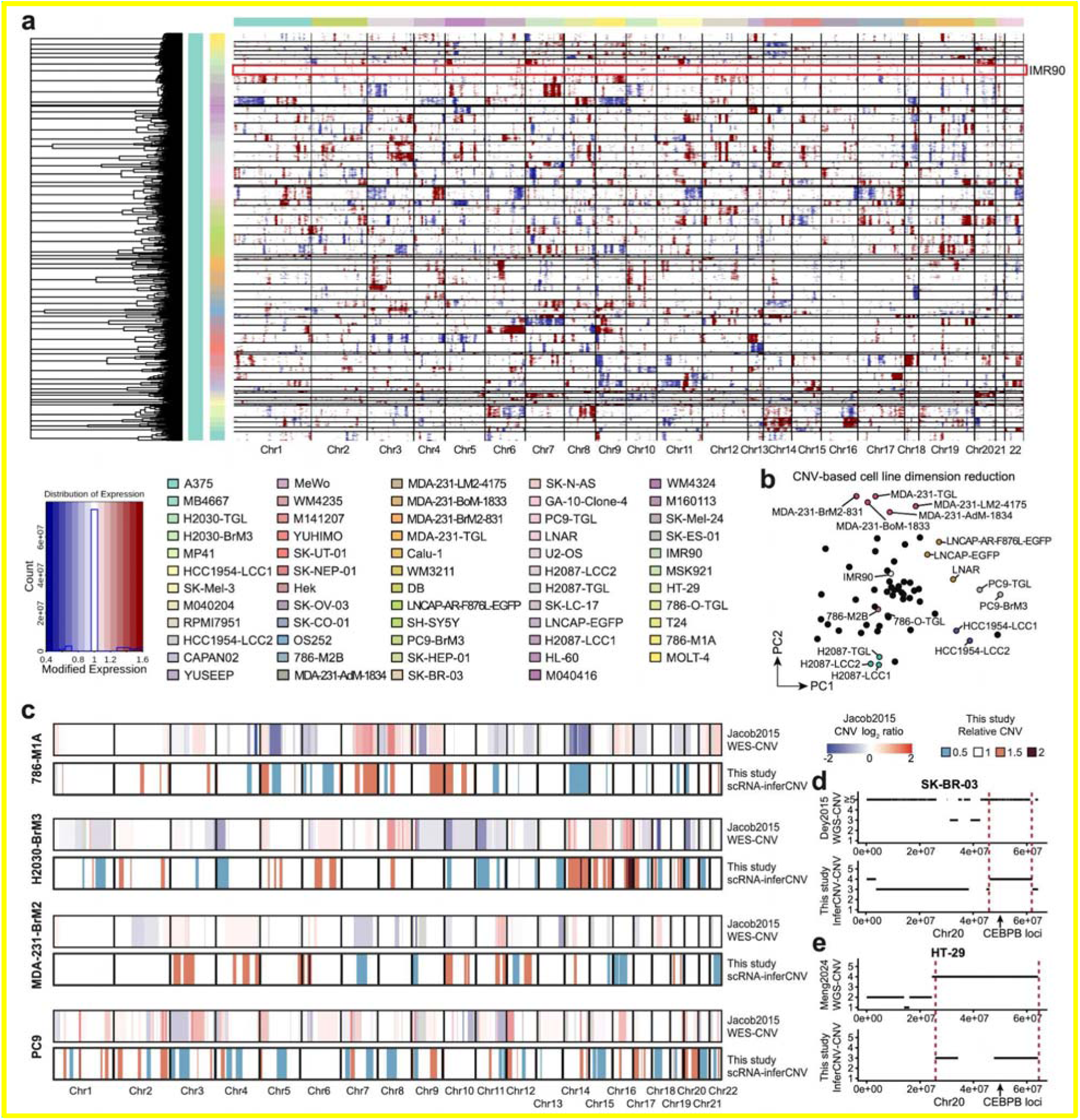
Pan-cancer cell line copy number variation inference by InferCNV. **a,** Heatmap showing smoothed RNA gene expression–based CNV patterns across meta-cells of pan-cancer cell lines. The karyotypically normal diploid cell line IMR90 is highlighted. Hierarchical clustering of CNV patterns was performed within each cell line, and no statistically significant subclones were identified. **b,** PCA plot showing the distribution of cell lines along the first two principal components based on inferred gene-level CNV profiles. Cell lines with close genetic evolutionary proximity, as well as IMR90, are highlighted. **c,** Comparison of genome-wide CNV profiles inferred from scRNA-seq in this study with published WES-derived CNV profiles from Jacob et al.^44^ for 786-M1A, H2030-BrM3, MDA231-BrM2, and PC9. **d,** Comparison of chr20 amplifications in SK-BR-03 between published WGS-derived CNV data^46^ and scRNA-seq-based inferCNV from this study. **e,** Comparison of chr20 CNV profiles in HT-29 between published WGS-derived CNV data^45^ and scRNA-seq-based inferCNV from this study.

To investigate the CNV landscape in the EMT process, we analyzed the significance of gene-level CNV-EMT pseudotime associations along genomic coordinates (**Fig. 3a**). In total, we identified 107 genes whose CNVs were significantly associated with pan-cancer EMT status, notably with exclusively negative correlations, indicating that amplification of these loci aligns with an epithelial cell state (**Fig. 3c**). As expected for genes affected by regional genetic alterations, these genes were concentrated in two specific genomic regions on chromosomes 11 and 20, respectively, which we refer to here as genomic hotspot loci (**Fig. 3c, Extended Data Fig. 6a-h**). Because ATAC chromatin accessibility reflects local DNA abundance, we used ATAC signal as an orthogonal proxy to validate transcriptome-derived CNV–EMT associations (**Methods, Extended Data Fig. 6a-h**). This approach highlighted 65 genes on chromosome 11 (*e.g., PPFIA1, CCND, LRP5, NADSYN1, PPP1CA*) and 8 genes on chromosome 20 (*e.g., CEBPB, B4GALT5, UBE2V1*), where amplification correlated negatively with EMT status (**Fig. 3c**). Amplification of these regions occurred predominantly in cell lines within the epithelial states, whereas no amplification or deletion was detected in mesenchymal states (**Fig. 3d**). The finding is consistent with our prior analysis of differentially accessible ATAC peaks across pan-cancer GRN clusters (**Fig. 2f**). For example, peaks specifically activated in GRN cluster 6 were enriched for genes located within these regions, such as *CCND1* and *PTPN2* in the chromosome 11 hotspot, and *CEBPB* and *UBE2V1* in the chromosome 20 hotspot (**Fig. 2f, Extended Data Fig. 6a-h**).

**Extended Data Fig. 6.**
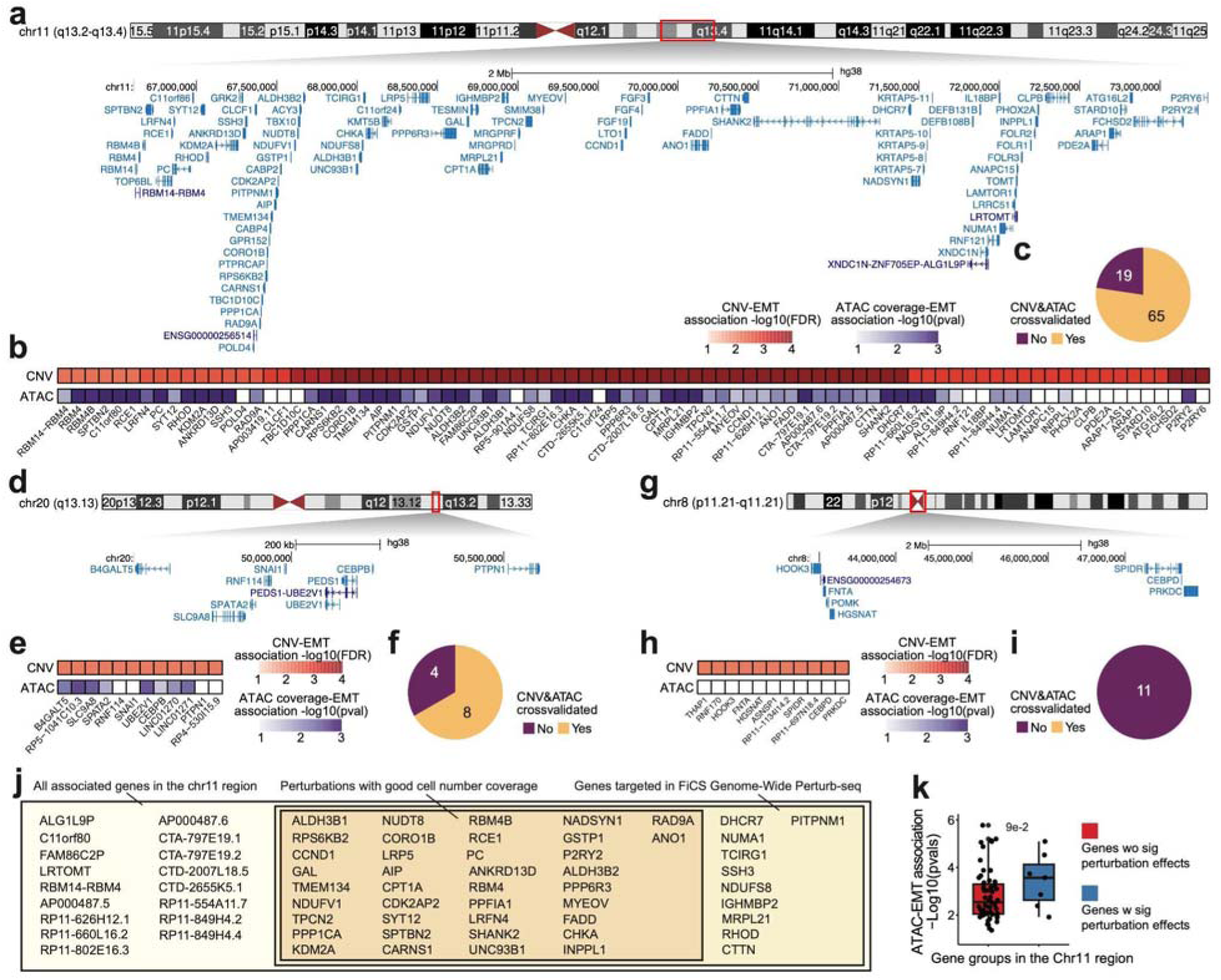
Characterization of the association between CNV of hotspot regions and EMT. **a,** Genome track displaying the gene distribution within the chromosome 11 hotspot region. The UCSC Genome Browser (hg38) was used for visualization, with RefSeq gene annotations shown. **b,** Heatmaps showing the statistical significance of gene expression–derived CNV–EMT pseudotime associations (top) and normalized ATAC coverage-based gene-EMT pseudotime associations (bottom) within the chromosome 11 hotspot region. White tiles indicate non-significant associations (Pearson correlation p ≥ 0.05). **c,** Pie chart showing the number of genes within the chromosome 11 hotspot region significantly associated with CNV-EMT pseudotime, and the subset validated by ATAC-EMT association. **d,** Genome track displaying the gene distribution within the chromosome 20 hotspot region. The UCSC Genome Browser (hg38) was used for visualization, with RefSeq gene annotations shown. **e,** Heatmaps showing the statistical significance of gene expression–derived CNV-EMT pseudotime associations (top) and normalized ATAC coverage-based gene-EMT pseudotime associations (bottom) within the chromosome 20 hotspot region. White tiles indicate non-significant associations (Pearson correlation p ≥ 0.05). **f,** Pie chart showing the number of genes within the chromosome 20 hotspot region significantly associated with CNV-EMT pseudotime, and the subset validated by ATAC-EMT association. **g,** Genome track displaying the gene distribution within the chromosome 8 region that failed ATAC validation. The UCSC Genome Browser (hg38) was used for visualization, with RefSeq gene annotations shown. **h,** Heatmaps showing the statistical significance of gene expression–derived CNV–EMT pseudotime associations (top) and normalized ATAC coverage–based gene–EMT pseudotime associations (bottom) within the chromosome 8 region. White tiles indicate non-significant associations (Pearson correlation p ≥ 0.05); no gene in this region exhibited a significant ATAC-EMT association. **i,** Pie chart showing the number of genes within the chromosome 8 region significantly associated with CNV–EMT pseudotime, and the subset validated by ATAC–EMT association. **j,** Venn diagram showing genes within the chromosome 11 hotspot region targeted in the FiCS Perturb-seq dataset. Genes not covered were primarily noncoding, and 10 perturbations were excluded due to low cell number coverage (n < 100). **k,** Box plot comparing the statistical significance of ATAC signal–EMT associations between genes with or without significant phenotypic shifts in the Perturb-seq data (no-shift group: n = 58; significant-shift group: n = 7). Boxes in box plots indicate the median and interquartile range (IQR), with whiskers indicating 1.5× IQR.

We next asked whether amplification of these genomic loci impacts drug response, given that cancer cell state is known to influence drug sensitivity^3,47^. Leveraging Cancer Cell Line Encyclopedia (CCLE)^48^, we associated CNV profiles of genes within these regions with IC50 values of anti-cancer drugs across large panels of cancer cell lines (**Fig. 3e**). By analyzing the number of genes whose amplification correlated with enhanced sensitivity (‘lower IC50’), we identified *Afatinib* as the top drug whose efficacy increased with amplification of most genes in both hotspot regions, followed by other EGFR and downstream inhibitors (**Fig. 3f**). This aligns with the established role of EGFR signaling as a key inducer of the EMT phenotype in cancer, and with prior evidence that EMT broadly confers resistance to EGFR pathway inhibitors^49–51^. Similarly, docetaxel, another drug previously reported to be resisted by EMT^52^, showed specific associations with a subset of genes in the chromosome 11 region (**Fig. 3f**).

Interestingly, *CEBPB*, a gene encoding a TF located in the chromosome 20 hotspot, was identified to be linked with the epithelial state of the EMT spectrum (**Fig. 3c**). When examining the association between inferred EMT pseudotime and the chromatin accessibility and gene expression-based CNV of *CEBPB*, it showed consistently increased ATAC accessibility and copy number amplification in epithelial-state lines, suggesting its hyperactivation as a potential mechanism preventing mesenchymal transition (**Fig. 3g**). As CEBPB was identified as a key TF within the pan-cancer GRN, we first used an in silico knockout strategy to examine how CEBPB ablation may affect cell state (**Methods**). In brief, SCENIC+ models gene expression based on the expression of inferred TF regulators and simulates perturbation-induced gene expression changes by setting CEBPB expression to zero. This process was iterated to simulate propagation of perturbation effects through the network. The simulated transcriptomes were then projected into the EMT PCA space to infer cell-state shifts. Consistent with our hypothesis, in silico CEBPB knockout induced a strong shift toward the mesenchymal state, specifically in epithelial-state cells (**Fig. 3h**). To test the generalizability of our analysis, we integrated paired CEBPB ChIP-seq^53^ and transcriptome profiles from external cancer cell lines (MCF-7, HeLa, A549, HepG2, and HCT116)^10^. By mapping these cell lines onto our inferred EMT trajectory (**Methods**), we estimated their EMT pseudotime and observed progressively reduced CEBPB binding at epithelial gene loci identified as CEBPB targets in our pan-cancer GRN, including KRT8 and KRT18^34^ (**Fig. 3i–j**). Finally, we leveraged CRISPRa Perturb-seq data from RPE1 epithelial cells with TF overactivation^54^ to test whether CEBPB activation is sufficient to enhance this regulatory program. Consistently, CEBPB activation led to increased activity of the CEBPB regulon identified from our pan-cancer atlas (**Fig. 3k**), including epithelial genes KRT18, KRT8, KRT7, KRT80, and JUP (**Fig. 3l**). Together, these orthogonal validations support a role for CEBPB in maintaining the epithelial identity of cancer cells.

In contrast, another hotspot region on chromosome 11 comprised a large cluster of genes but lacked major transcriptional regulators (**Extended Data Fig. 6a**). To distinguish potential drivers from passenger genes, we reanalyzed a published genome-wide CRISPRi Perturb-seq dataset^55^ that profiled epithelial cells with individual gene silenced, in which most genes within this region were perturbed (**Fig. 3m**, **Extended Data Fig. 6j**). To examine the potential contribution of these genes to EMT-associated state changes, we extracted top marker genes representing cell lines in GRN clusters 6 and 8 and defined them as epithelial and mesenchymal features, respectively. We then compared these phenotypic programs across individual gene perturbations relative to control cells. Mimicking the region-level negative CNV–EMT association, the aggregate perturbation effects showed an overall loss of epithelial features and activation of mesenchymal features (**Fig. 3n**). At the individual perturbation level, while most perturbations showed no observable effect (n=31) or only partial effects on the epithelial program (n=2), perturbation of LRP5, RBM4B, NADSYN1, SYT12, and TMEM134 (n=5) induced both significant activation of the mesenchymal program and downregulation of the epithelial program relative to NTC cells (**Fig. 3o**). Strong induction of representative mesenchymal markers, including FN1, CDH2, and MMP16, further supported causal roles of these genes in maintaining the epithelial state (**Fig. 3p**). Notably, in our discovery dataset, these genes whose perturbation significantly altered epithelial and/or mesenchymal features in the Perturb-seq validation analysis (n=7) exhibited stronger ATAC accessibility–EMT associations than the remaining genes in the region (**Fig. 3q, Extended Data Fig. 6k**).

In summary, our systematic pan-cancer CNV characterization highlights TF amplification, followed by their hyperactive transcriptional regulation, as a major mechanism shaping cancer cell states. CNV–EMT association analyses also indicate mechanistic heterogeneity: while *CEBPB* amplification reinforces epithelial identity via direct transcriptional regulation, regional amplifications also likely involve multiple genes with heterogeneous individual effects, whose combined dosage ultimately shapes the cell-state outcome. These distinct routes converge on cell-state modulation but also likely carry divergent implications for drug sensitivity.

### Subtype-resolved gene regulatory network in acral and cutaneous melanoma

We next assessed whether similar regulatory patterns are preserved in heterogeneous melanoma cells isolated from different patients, with a specific focus on acral melanoma. Acral melanoma (AM) is a rare melanoma subtype with distinct epidemiology and prognosis^56^, but remains poorly characterized at the molecular level. To investigate its molecular signatures, we included diverse melanoma cell lines in our pan-cancer cell line cohort, which can be anatomically classified as uveal melanoma (n=1), acral melanoma (AM) (n=10 after quality control), and cutaneous melanoma (CM) cell lines (n=5) (**Fig. 1b**). Consistent with their melanocyte origin, 14 of 15 cell lines showed highly similar gene-regulatory features within the pan-cancer gene regulatory network, forming cluster 5 (**Fig. 2b**) and exhibiting stable lineage identity with high MITF and SOX10 activity (**Fig. 2c**).

To examine the molecular differences between acral melanoma (AM) and cutaneous melanoma (CM), we performed Orthogonal Non-negative Matrix Factorization (oNMF)^57^ in combination with differential expression analysis^58^ on the dataset (**Methods**). Extensive benchmarking supported the robustness of the decomposed gene programs (**Extended Data Fig. 7a-d**). Differentially expressed genes between AM and CM recapitulated their most recognized divergence: CM displayed transcriptomic features of UV exposure, whereas AM, typically arising from sun-shielded skin, lacked such signatures (**Extended Data Fig. 7e-f**). Among the 15 oNMF-derived programs, 11 programs were strongly activated in individual cell lines (**Fig. 4a**), reflecting pronounced inter-cell line heterogeneity (**Fig. 1c-d**). Major phenotypic states of melanoma^59^, irrespective of subtype, were captured in this cohort: Program 0, enriched for pigmentation genes such as *MITF*, *TYR*, *OCA2*, *DCT*, strongly associated with the melanocytic differentiation but inversely correlated with melanoma dedifferentiation (**Fig. 4b**), whereas Program 1, enriched for neural crest–associated genes such as *NGFR*, *CDH*2, *GDNF, GFRA1*, mirrored the predefined neural crest-like state (**Fig. 4b**). Consistent with these associations, SK-Mel-3, the melanoma cell line with the lowest Program 0 score (**Fig. 4b**), exhibited the strongest expression of the undifferentiated signature and was the only melanoma cell line clustered within the cross-lineage mesenchymal cluster (cluster 8) in the pan-cancer GRN (**Fig. 2b**).

**Fig. 4.**
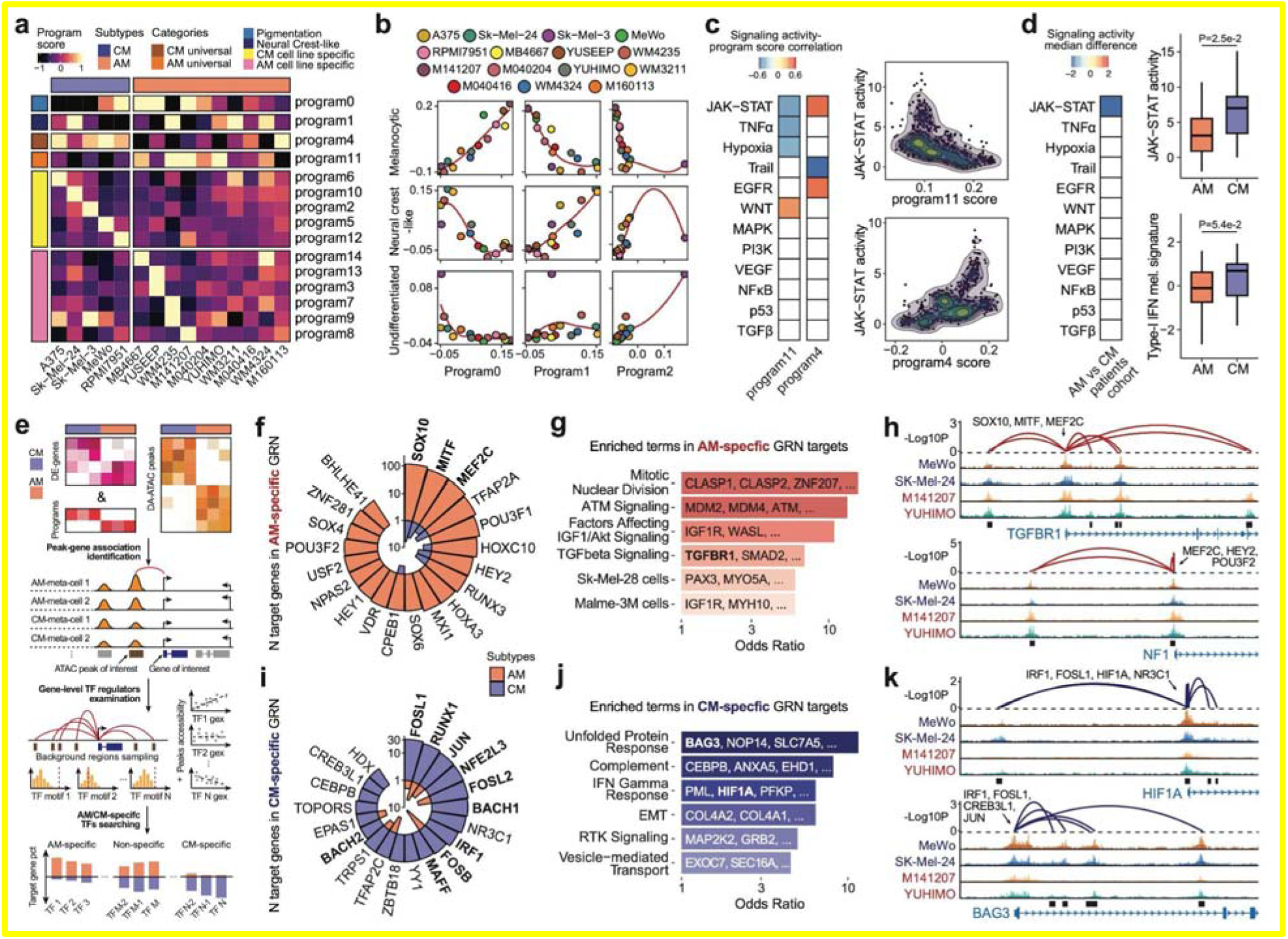
Integrated analyses define distinct gene regulatory principles across melanoma subtypes despite phenotypic diversity. **a,** Heatmap showing the mean scores of orthogonal gene programs deconvolved from skin cancer cell lines. **b,** Scatter plots showing associations between established melanoma phenotypic signatures and oNMF-derived gene programs. In each plot, one axis shows a gene program score and the other a melanoma signature score, and each dot represents the mean score for a cell line. Colors denote individual melanoma cell lines (as in Fig. 4a), and error bars indicate the standard deviation of scores across meta-cells within each cell line. Numerical axis scales are omitted to emphasize relative trends. **c,** Heatmaps of correlations between signaling activities inferred from meta-cell transcriptomes and AM/CM universal program scores across skin cancer cell lines (left panel). Scatter plots with kernel density overlays showing JAK–STAT activity versus program scores across meta-cells (right panel). **d,** Heatmap showing differences in mean AM/CM universal program scores across patients in the validation cohort (left panel). Only signaling activities with significant subtype differences are shown. Box plots of JAK–STAT activity scores and type-I IFN melanoma signature scores by subtype (right panel). AM patients n=42, CM patients n=15. **e,** Schematic of melanoma subtype–specific GRN construction and identification of TF regulators. **f,** Rose plot displaying AM-specific TFs and the number of their target genes within each DEG-refined subtype-specific program. TFs emphasized in the main text are bolded. **g,** Bar plot showing enrichment of functional terms among AM-specific TF target genes; representative genes within each term are labeled. **h,** Representative genome tracks showing pseudobulk-normalized ATAC signal coverage around TGFBR1 and NF1, genes universally upregulated in AM. Expression-associated peaks are boxed; CRE–gene linkages are shown as arches; significant TF regulators of each CRE–gene pair are indicated. Example CM cell lines are labeled in blue and AM cell lines in red. **i,** Rose plot displaying CM-specific TFs and the number of their target genes within each DEG-refined, subtype-specific program. TFs emphasized in the main text are bolded. **j,** Bar plot showing enrichment of functional terms among CM-specific TF target genes; representative genes within each term are labeled. **k,** Representative genome tracks showing pseudobulk-normalized ATAC signal coverage around HIF1A and BAG3, genes universally upregulated in CM. Expression-associated peaks are boxed; CRE–gene linkages are shown as arches; significant TF regulators of each CRE–gene pair are indicated. Example CM cell lines are labeled in navy and AM cell lines in firebrick. Boxes in box plots indicate the median and interquartile range (IQR), with whiskers indicating 1.5× IQR. Panel d uses an external patient validation cohort (bulk RNA-seq). Panel b benchmarks our cell-line gene programs against published melanoma phenotypic signatures (Tsoi et al. and Wouters et al.); panel e is a schematic. All other panels show analyses generated in this study.

**Extended Data Fig. 7.**
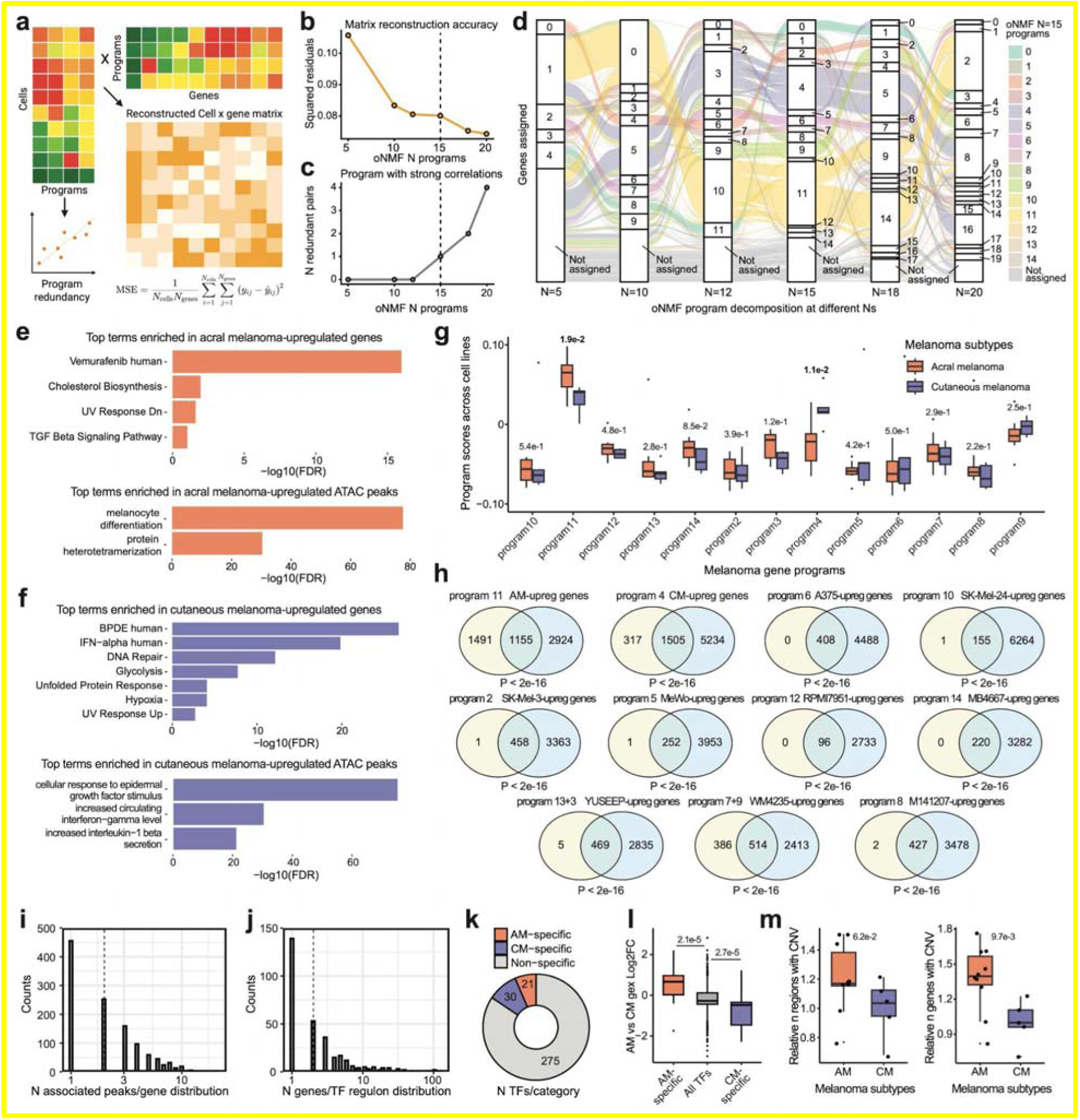
Characterization of Melanoma subtype-specific gene-regulatory features. **a,** Schematic showing evaluation of oNMF decompositions by matrix reconstruction accuracy and program redundancy. **b,** Mean squared reconstruction residuals across different oNMF program numbers, showing diminishing improvement after approximately 10 programs; dashed line indicates the selected N=15. **c,** Number of redundant program pairs, defined by Pearson correlation >0.65 between program scores across melanoma cells, across different program numbers; dashed line indicates N=15. **d,** Sankey diagram showing conservation of gene membership across decompositions, with genes colored by their program identity in the selected N=15 decomposition. **e,** Bar plots showing the top functional terms significantly enriched among upregulated genes (top) and ATAC peaks (bottom) in AM relative to CM. EnrichR and GREAT were used for gene-based and chromatin accessibility–based enrichment analyses, respectively. **f,** Bar plots showing the top functional terms significantly enriched among upregulated genes (top) and ATAC peaks (bottom) in CM relative to AM. EnrichR and GREAT were used for gene-based and chromatin accessibility–based enrichment analyses, respectively. **g,** Box plot comparing pseudobulk gene program score distributions between AM and CM cell lines. P-values were calculated using two-sided Wilcoxon tests. **h,** Venn diagrams showing overlaps between oNMF-derived orthogonal gene programs and significantly differentially expressed genes identified by DESeq2. Overlapping genes were considered DEG-refined program members. P-values were calculated using Fisher’s exact tests. **i,** Histogram showing the distribution of the number of ATAC peaks associated with downstream genes. The dashed line indicates the median (2). **j,** Histogram showing the distribution of the number of target genes per TF. The dashed line indicates the median (2). **k,** Donut chart showing the number of TFs assigned to different melanoma regulatory categories. **l,** Box plot showing the AM versus CM relative expression changes of TFs identified as AM-specific (n = 21) or CM-specific (n = 30) through GRN construction. The full list of TFs examined in FigR was used as background (n = 1141). Two-sided p-values from Tukey’s tests following one-way ANOVA are reported. **m,** Box plots showing the distributions of the number of genomic regions with CNV (left) and the number of genes with CNV (right) between AM and CM cell lines (AM, n = 10; CM, n = 5). Two-sided p-values from Wilcoxon tests are reported. Boxes in box plots indicate the median and interquartile range (IQR), with whiskers indicating 1.5× IQR.

Importantly, we observed that Program 4 and Program 11 exhibited consistent activation in either CM or AM cells (AM universal genes n=1,155, CM universal genes n=1,505), revealing the existence of subtype-specific transcriptional modules distinguishing cutaneous and acral melanoma (**Extended Data Fig. 7g**). These gene programs also significantly overlapped with differentially expressed genes between AM and CM (**Extended Data Fig. 7h**). By examining signaling activities across melanoma cell lines, although several pathway activities correlated with one of the DEG-refined subtype-universal gene programs, JAK–STAT signaling uniquely showed significant, but opposite associations in Program 4 and Program 11 (**Fig. 4c**). Specifically, we found JAK-STAT signaling negatively correlates with the AM-universal program (R = -0.52, FDR = 3.68e-72) and positively with the CM-universal program (R = 0.53, FDR = 1.34e-76), which indicates an attenuated inflammatory phenotype in AM. Scoring bulk RNA-seq from a patient cohort containing both AM and CM^60^ further validated this pattern: only JAK-STAT activity exhibited a significant reduction in AM relative to CM (**Fig. 4d**). Moreover, the score of a Type-I Interferon (IFN) signature, defining a subset of melanoma cells from patient samples^60^, was consistently lower in AM (**Fig. 4d**), indicating a universal inflammatory dampened state in this subtype.

We next explored the regulatory logic underlying these subtype distinctions by integrating refined AM- and CM-universal gene programs with subtype-specific differentially accessible ATAC peaks (AM-specific peaks, n=63,510; CM-specific peaks, n=40,862). This enables the construction of melanoma subtype-specific CRE–gene linkages and the identification of potential master TF regulators (**Fig. 4e, Extended Data Fig. 7i-j, Methods**). Among 326 TFs identified to regulate at least 1 gene in melanoma, 21 were classified as AM-specific and 30 as CM-specific (**Extended Data Fig. 7k-l**).

Consistent with our prior characterization, melanocytic TFs predominated in the AM-specific GRN (**Fig. 4f**), which are known to suppress inflammatory gene programs^61,62^. Target genes of these TFs were enriched for the IGF pathway (**Fig. 4g**), previously implicated in specifically promoting AM pathogenesis^63^. Enrichment of ATM signaling further reflected pervasive genomic instability in AM, in contrast to the UV radiation-induced point mutations characteristic of CM^64–66^, consistent with our comparison of CNV patterns across melanoma subtypes (**Extended Data Fig. 7m**). The convergence of these features with transcriptomic signatures of MITF-dependent melanocytic cell lines such as SK-Mel-28^67^ and Malme-3M^68^ (**Fig. 4g**) further supports the role of these TFs in shaping the gene-regulatory landscape of AM. For instance, *TGFBR1*, encoding the key receptor of the TGF-β pathway known to suppress intracellular inflammatory activity^69^, exhibited AM-specific upregulation through melanocytic TFs-associated CREs; *NF1*, a negative regulator of MAPK signaling, was activated through MEF2C-associated CREs in AM (**Fig. 4h**), adding a transcriptional regulatory layer to the mutual suppression between melanocytic and MAPK activities in melanoma^70^.

In contrast, the gene regulatory landscape of CM was characterized by extensive involvement of inflammatory TFs (**Fig. 4i**), dominated by AP-1 factors (*FOSL1, JUN, FOSL2, FOSB*), *RUNX1*, and CNC/bZIP factors (*NFE2L3, BACH1, MAFF, BACH2*) that share core motifs with AP-1. Notably, IRF1, an essential transcriptional effector of IFN stimulation and JAK-STAT signaling^71^, also emerged as one of the top CM-specific regulators (**Fig. 4i**), reinforcing the predominance of inflammatory transcriptional activity. Consistently, CM-specific target genes were enriched in inflammation-related functions, including the unfolded protein response and IFN responses (**Fig. 4j**). For example, the CRE associated with *HIF1A* and *BAG3* displayed increased chromatin accessibility in several CM cell lines and were enriched for AP-1 and IRF1 motifs (**Fig. 4k**). Together, these findings suggest that acral melanoma exhibits a universal inflammation suppressive phenotype, while cutaneous melanoma is represented by an inflamed phenotypic and gene regulatory landscape, between which JAK-STAT, along with downstream transcriptional response, might serve as a key player.

### From cancer-intrinsic programs to tumor microenvironment remodeling and prediction of immunotherapy response

Extensive studies have highlighted the central role of cancer cells in reprogramming their surrounding cellular niche through complex intercellular communication^72–74^. These adaptive changes in the tumor microenvironment (TME) can, in turn, substantially affect cancer cell behavior and therapeutic responses^75^. Given that inflammatory programs involve both intracellular and intercellular activities, we hypothesized that these key melanoma-intrinsic programs observed in vitro may be associated with distinct features of the original tumor microenvironment. To test this hypothesis, we explored bulk RNA-seq profiles from the TCGA SKCM cohort, in which the measured transcriptomes reflect both tumor-intrinsic and infiltrating TME compartments. We assessed associations between melanoma program scores, representing tumor-intrinsic states, and deconvolved TME cell-type scores, representing the microenvironmental compartment. This analysis revealed broad negative correlations between the AM-specific Program 11 and multiple immune-supportive populations across patients, including cytotoxic NK cells (R = −0.39, FDR = 1.5e−5), CD8⁺ T effector memory cells (R = −0.25, FDR = 8.6e−3), CD4⁺ Th1 cells (R = −0.63, FDR = 8.0e−16), antigen-presenting dendritic cells (R = −0.39, FDR = 1.5e−5), and endothelial cells (R = −0.43, FDR = 1.1e−6), which facilitate tumor immune infiltration (**Fig. 5b, c**)^76^. In contrast, Tregs, a major immunosuppressive cell type in the TME, showed a positive association with Program 11 (R = 0.32, FDR = 4.1e−4). These patterns are concordant with a previous single-cell study reporting reduced effector T cell infiltration and increased Tregs among tumor-infiltrating lymphocytes from AM patients^77^. These associations suggest that melanoma subtype-specific cancer-intrinsic programs detected in vitro are linked to distinct TME features in patient tumors. Moreover, after stratifying TCGA SKCM samples by AM- and CM-specific program scores (Fig. 5c), we found significantly lower predicted MHC-I neoantigen burden^78^ in AM-like samples (Program 11-high, Program 4-low) than in CM-like samples (Program 11-low, Program 4-high). This is consistent with the high UV-driven tumor mutation burden of CM^79^, and the relatively immunosuppressive TME landscape associated with AM.

**Fig. 5.**
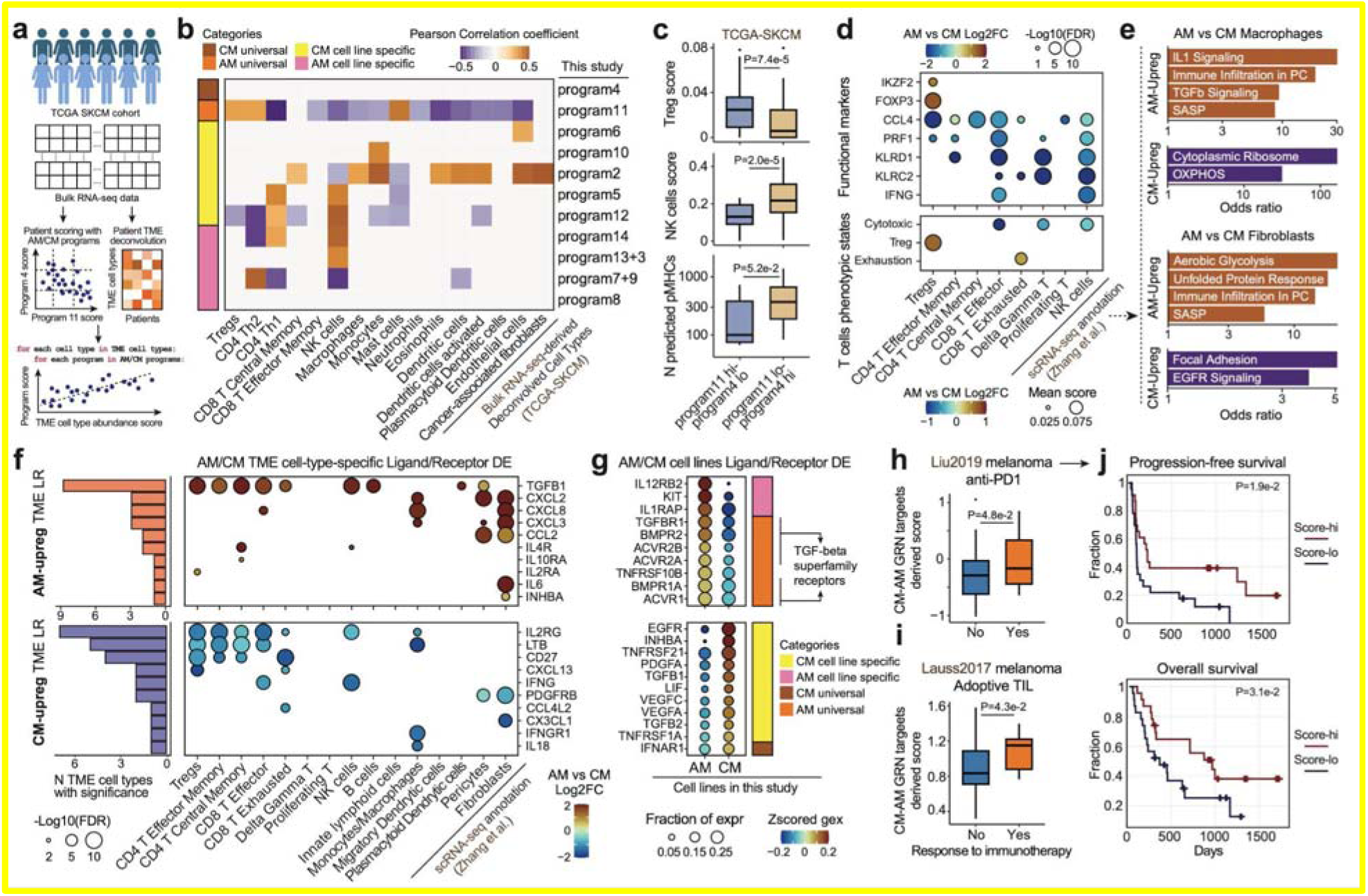
Subtype-resolved gene regulatory signatures from melanoma cell lines map TME states and immunotherapy sensitivity. **a,** Schematic illustrating the analytical strategy used to process RNA-seq profiles from the TCGA SKCM cohort. **b,** Heatmap showing the correlation between melanoma subtype–specific gene program scores and deconvolved TME cell-type scores. Categories of gene programs are defined in Fig. 4a. **c,** Boxplots showing differences in Treg cell scores (top), NK cell scores (middle), and the number of peptides predicted to bind MHC (bottom) between TCGA SKCM patients exhibiting strong AM or CM transcriptomic features. Program 11 hi–program4 lo patients, n=46; Program 11 lo–program 4 hi patients, n=56. In the pMHC panel, only samples with whole genome-sequencing data were used: program11 hi–program4 lo patients, n=6; program 11 lo–program 4 hi patients, n=16. **d,** Dot heatmap showing relative log2 fold changes of representative functional markers (top) of T cells and T cells immune signature scores characterized from published scRNA-seq datasets of patient samples. **e,** Bar plots showing the top enriched functional terms among DEGs in macrophages (top) and fibroblasts (bottom) from AM patients compared with CM patients. **f,** Dot heatmap showing relative log2 fold changes of ligand/receptor genes between AM and CM TME cell types. The top panel shows ligand/receptor genes upregulated in AM, and the bottom panel shows those upregulated in CM. Genes are ordered by the number of cell types showing significant differences, with the counts displayed in the left columns. **g,** Dot heatmap showing z-scored expression levels of significantly differentially expressed ligand/receptor genes between AM and CM cell lines that also belong to one of the identified gene programs. **h,** Boxplots comparing biomarker scores computed using CM-specific versus AM-specific target genes from pre-treatment patient RNA-seq data between responders and non-responders under anti–PD-1 therapy (n=17 responders; n=29 non-responders). The P-value from the two-sided Wilcoxon test was displayed. **i,** Boxplots comparing biomarker scores computed using CM-specific versus AM-specific target genes from pre-treatment patient RNA-seq data between responders and non-responders under adoptive tumor-infiltrating lymphocyte therapy (n=10 responders; n=15 non-responders). The P-value from the two-sided Wilcoxon test was displayed. **j,** Kaplan–Meier survival curves showing progression-free survival (top) and overall survival (bottom) of patients with high versus low biomarker scores computed using CM-specific versus AM-specific target genes (n=23 for high-score; n=23 for low-score). The two-sided Wald p-value for the signature term from the univariate Cox regression was reported. Boxes in box plots indicate the median and interquartile range (IQR), with whiskers indicating 1.5× IQR. Panel a is a schematic of the TCGA-SKCM analysis. Panel a is a schematic of the TCGA-SKCM analysis. External patient data are used in panels b-c (TCGA-SKCM), d-f (published patient single-cell RNA-seq), and h-j (immunotherapy patient cohorts); panel g shows our cell-line data.

To evaluate potential confounding of bulk melanoma subtype-associated program scores by tumor purity or TME composition, we performed additional analyses using patient melanoma single-cell RNA-seq data and TCGA-SKCM cohort (**Extended Data Fig. 8**). Both Programs 4 and 11 were strongly enriched in melanoma cells relative to non-malignant TME compartments (**Extended Data Fig. 8a**). TCGA-SKCM samples generally showed high tumor purity^81^ (**Extended Data Fig. 8b**), and the bulk scores of these cancer-intrinsic programs were positively associated with estimated tumor purity (**Extended Data Fig. 8c**). Moreover, residual Program 11 gene scores showed no change in TME cell types between AM and CM patients (**Extended Data Fig. 8d**), supporting the interpretation of Program 11 as a melanoma-cell-derived program.

**Extended Data Fig. 8.**
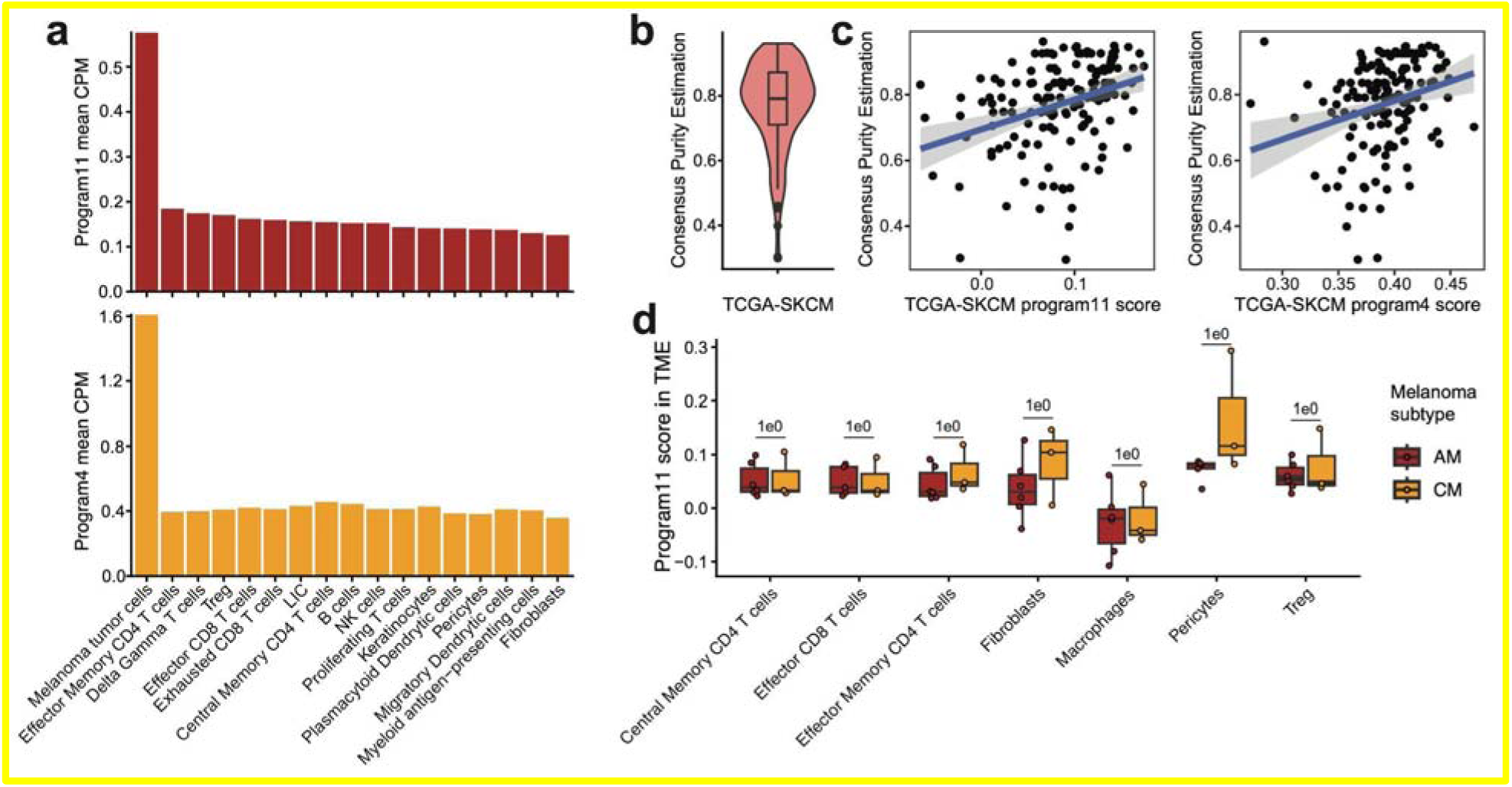
Evaluation of tumor-cell specificity and TME spill-over for melanoma subtype-associated programs. **a,** Mean CPM expression of program 11 and program 4 genes across melanoma tumor cells and non-malignant TME cell types in patient melanoma single-cell RNA-seq data. **b,** Distribution of consensus tumor purity estimates across TCGA-SKCM bulk RNA-seq samples. **c,** Association between consensus tumor purity estimates and bulk program 11 or program 4 scores in TCGA-SKCM samples. **d,** Comparison of program 11 scores in non-malignant TME cell types between AM and CM patient single-cell RNA-seq samples.

To further validate the suppressive microenvironment of AM at a molecular phenotype level, we further analyzed published single-cell RNA-seq of TME of both AM and CM patient samples^60^ (**Extended Data Fig. 9a-c**). Across cell types, T cell populations exhibited the most pronounced transcriptomic changes (**Extended Data Fig. 9d**). By comparing the expression of functional markers in T cell populations between AM and CM patients, we observed a global attenuation of antitumor immunity in AM (**Fig. 5d**). For example, Tregs in AM showed higher expression of IKZF2 and FOXP3, consistent with enhanced suppressive function. In contrast, PRF1 and IFNG, which encode core cytotoxic effector molecules and cytokines, showed markedly lower expression in CD8 effector T cells and NK cells in AM. By further quantifying phenotypic changes using recurrent T cell state signatures^82^, we found that Tregs and exhausted CD8 T cells from AM patients showed increased immunosuppressive/exhaustion-associated programs, whereas CD8 effector T cells, γδ T cells, and NK cells showed reduced cytotoxic programs (**Fig. 5d**). Other major cellular lineages in the TME also exhibited convergent immunosuppressive features. For example, macrophages from AM patients showed enrichment of IL1 signaling and senescence-associated secretory phenotype (SASP) programs, consistent with a tumor-associated macrophage (TAM)-like phenotype that may contribute to an immunosuppressive tumor microenvironment^83^. Fibroblasts from AM patients also showed an increased SASP secretion program, together with reduced focal adhesion and EGFR signaling, suggesting a shift toward an immunosuppressive cancer-associated fibroblast (CAF)-like state rather than a classic CAF phenotype^84^. Overall, these cell-type-specific transcriptomic profiles of the AM TME complement our melanoma gene program–TME composition inference and further substantiate a suppressive microenvironment aligned with the uninflamed melanoma cell state as a defining characteristic of AM.

**Extended Data Fig. 9.**
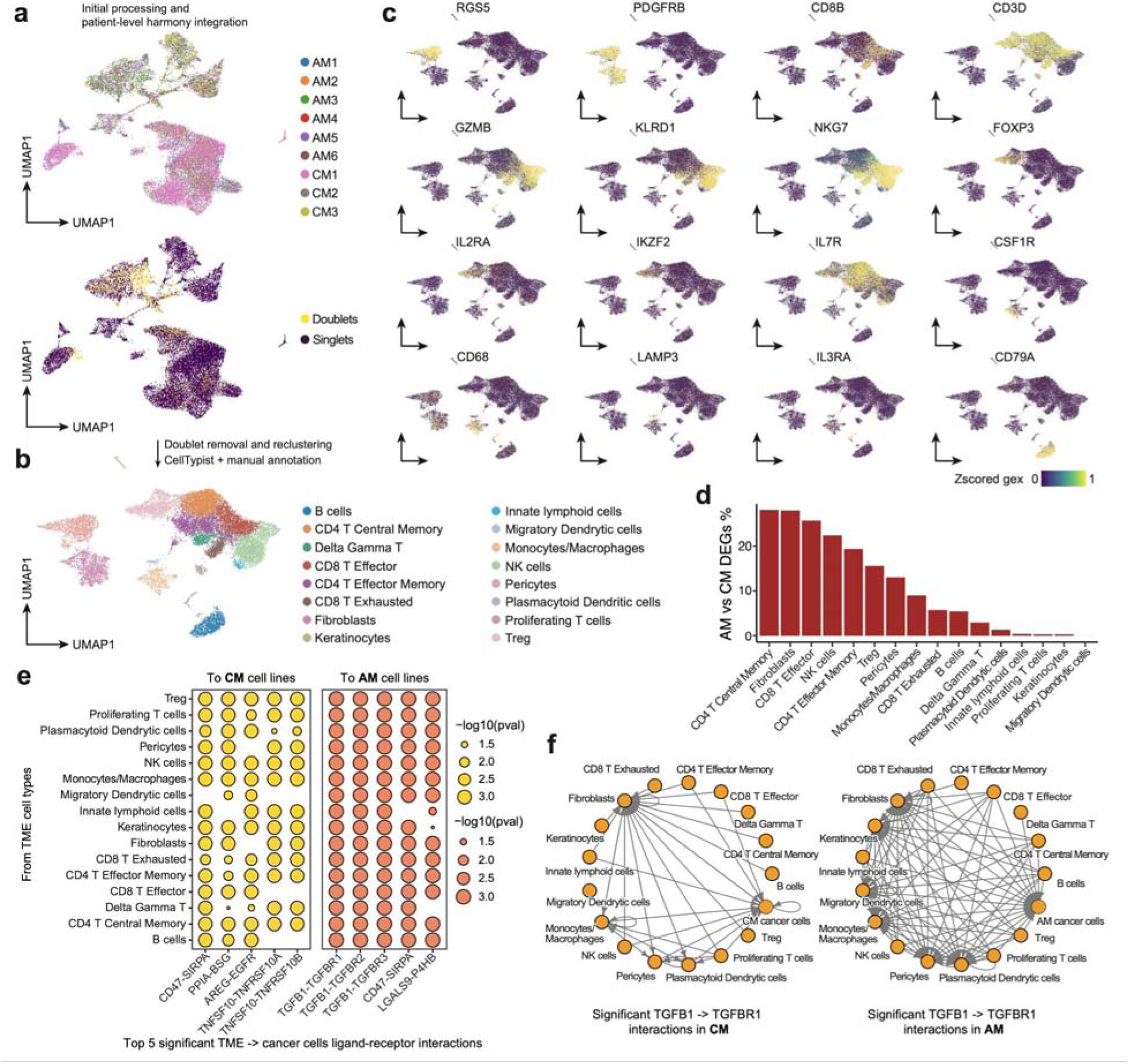
Patients’ TME single-cell RNA-seq data reprocessing and examinations of melanoma subtype-associated changes. **a,** First-round patient-level Harmony-integrated UMAP of TME single-cell RNA-seq from AM and CM patients. Data was reprocessed from^60^. Cells are colored by patient identity (top) and computationally identified doublets (bottom). **b,** Final patient-level Harmony-integrated UMAP of TME single-cell RNA-seq from AM and CM patients. Cells are colored by cell-type labels annotated using CellTypist^92^ and manual curation. **c,** UMAP colored by representative marker genes of major TME cell types. **d,** Bar plot showing the percentage of significantly differentially expressed genes for each TME cell type between AM and CM patients. **e,** Dot heatmaps showing the top five ligand–receptor pairs with the highest number of significant paired activations between TME cell types (ligand genes) and CM or AM cell lines (receptor genes), identified using CellPhoneDB^85^. **f,** Network diagrams showing all significant TGFB1–TGFBR1 paired activations across TME cell types and cancer cell lines in the CM (left) or AM (right) subtype, identified using CellPhoneDB.

To uncover candidate mediators linking cancer cells and TME, we compared ligand/receptor expression between AM and CM across both patients’ TME and melanoma cell lines (**Fig. 5e, f**). In TME, TGFB1 emerged as the most consistently upregulated ligand/receptor gene in AM, activated across 9 TME cell types, whereas others were mostly cell-type-specific (**Fig. 5e**). Across melanoma cell lines, TGF-β superfamily receptors exhibited universally activated expression in AM versus CM, and were identified as members of the AM universal gene program (Program 11). Among them, TGFBR1, encoding the receptor of TGF-β1, and downstream effectors such as SMAD2, were actively induced through the AM-specific GRN (**Fig. 4j-k**). Analysis of paired ligand/receptor expressions across compartments via CellPhoneDB^85^ showed a consistent trend, highlighting the potential central role of TGF-β in reprogramming TME and AM (**Extended Data Fig. 9e-f**). Consistently, TGF-β signaling is well established as a major suppressive axis in the TME: it inhibits cytotoxic programs of CD8⁺ T cells and NK cells, impedes CD4⁺ Th1 differentiation, downregulates MHC-II in dendritic cells, and enhances Treg activity through FOXP3 induction^86^.

Conversely, in CM, the most consistently upregulated ligands/receptors within the TME were IL2RG, LTB, and CD27 across T-cell populations, highlighting an activated antitumor milieu. The cancer cell compartment exhibited a parallel interactome landscape: while most cytokines upregulated in CM were cell-line-specific, particularly within the highly dedifferentiated SK-Mel-3, IFNAR1, which encodes the Type-I IFN receptor, was the only significantly upregulated ligand/receptor within the CM universal gene program, consistent with CM’s inflamed TME, elevated intracellular JAK-STAT signaling activity, and an inflammatory GRN (**Fig. 4i–k**).

Finally, because TME is a strong predictor of immunotherapy outcomes, we evaluated whether melanoma subtype-specific gene-regulatory features could predict immunotherapy response. Using the Cancer Immunology Data Engine (CIDE)^87^, which integrates pretreatment RNA-seq profiles and clinical metadata from immunotherapy cohorts, we tested AM- and CM-specific gene signatures as biomarkers of therapeutic response. Although the full AM and CM universal programs did not distinguish responders from non-responders (**Extended Data Fig. 10a-b**), incorporating gene-regulatory information by refining these programs to target genes of AM- and CM-specific TFs (**Fig. 4e-k**) markedly improved performance. These refined GRN-based scores significantly separated responders from non-responders from the same cohorts (**Fig. 5h-i**). Moreover, because anti-CTLA-4 and anti-PD-1 therapies are standard immunotherapeutic approaches for melanoma^88^, we further examined the prognostic potential of our melanoma subtype signatures (**Extended Data Fig. 10c**). Consistently, full AM and CM universal programs did not distinguish treated patients’ prognosis (**Extended Data Fig. 10d**). In anti-CTLA-4-naive patients treated with anti-PD-1, none of the pre-established immune signatures significantly predicted prognosis by univariate Cox regression, with the strongest-performing signature, IMPRES^89^, showing two-sided Wald P values of 0.111 for PFS and 0.122 for OS (**Extended Data Fig. 10e,f**). In contrast, the GRN-based scores specifically stratified prognosis in anti-CTLA-4-pretreated patients subsequently treated with anti-PD-1, achieving predictive performance comparable to established immune signatures and stronger than tumor mutation burden (**Fig. 5i**). Validation in additional cohorts^90,91^ further supported the generalizability of this finding (**Extended Data Fig. 10h-k**). This association remained significant for overall survival in multivariable Cox regression adjusting for age and sex (**Extended Data Fig. 10k**). Together, these results underscore the close association between melanoma-intrinsic regulatory logic and therapy outcomes, particularly in the anti-CTLA-4 to anti-PD-1 treatment sequence.

**Extended Data Fig. 10.**
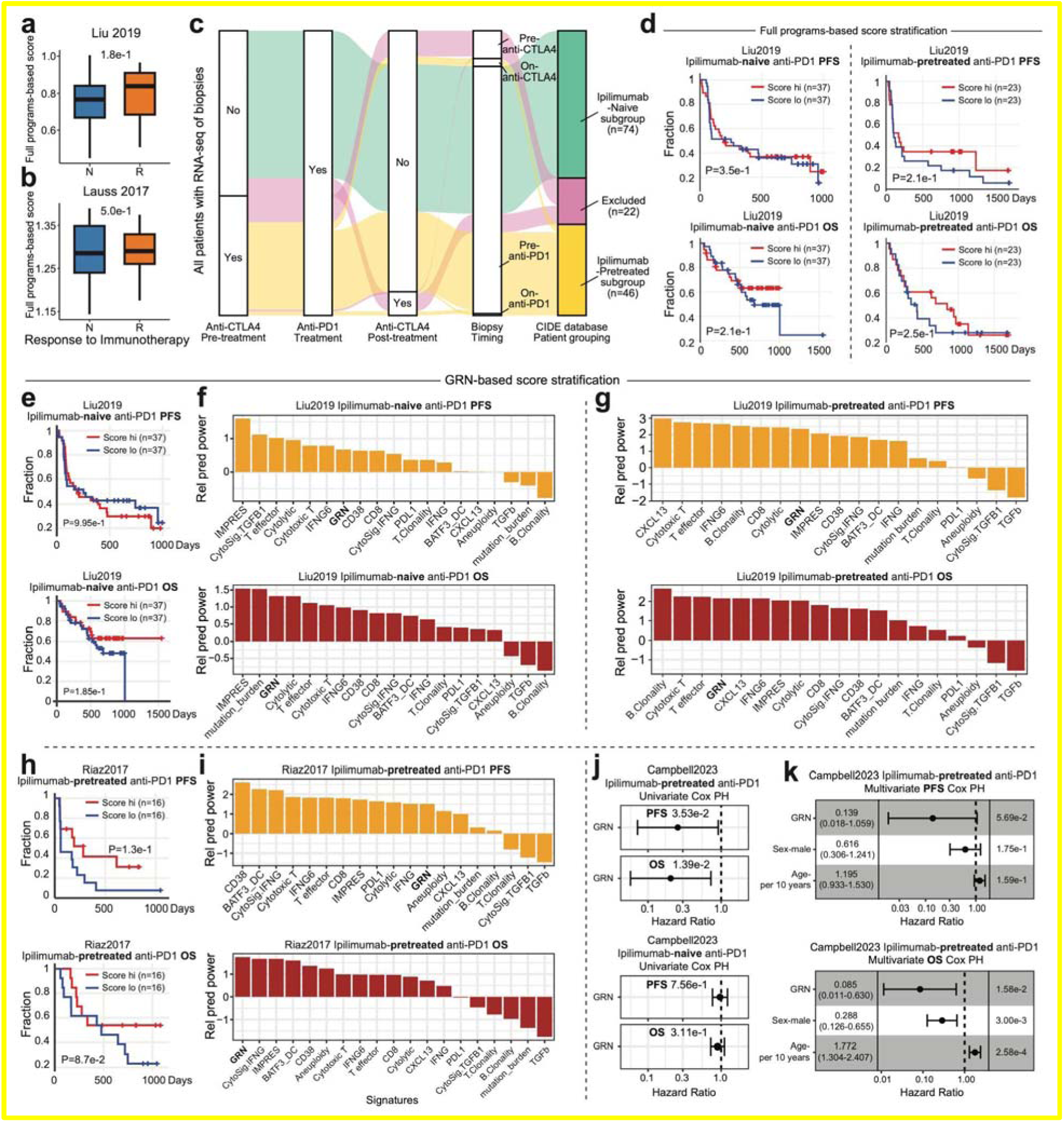
Comparative analyses of the power of gene signatures in immunotherapy patient cohorts. **a,** Box plot comparing scores of anti-PD-1 responder and non-responder from Liu2019^93^, calculated as full-gene Program 4 (CM-universal) minus full-gene Program 11 (AM-universal). P-value, two-sided Wilcoxon test. **b,** Box plot comparing scores of adoptive TIL therapy responder and non-responder from^94^, calculated as full-gene Program 4 minus full-gene Program. n = 10 responders; n = 15 non-responders. P-value, two-sided Wilcoxon test. **c,** Sankey diagram classifying Liu2019 anti-PD-1-treated melanoma patients by prior anti-CTLA-4 treatment, post-treatment anti-CTLA-4 exposure, and biopsy timing. **d,** Kaplan–Meier curves showing PFS and OS in Liu2019 anti-PD-1-treated patients stratified by ipilimumab-naive (left) or ipilimumab-pretreated (right) status and full-program scores. P-value, two-sided Wald test for the signature term from univariate Cox regression. **e,** Kaplan–Meier curves showing PFS (top) and OS (bottom) in ipilimumab-naive, anti-PD-1-treated Liu2019 patients stratified by GRN scores. P-value, two-sided Wald test for the signature term from univariate Cox regression. **f,** Bar plots showing the relative predictive power of gene signatures for PFS (top) and OS (bottom) in ipilimumab-naive, anti-PD-1-treated Liu2019 patients. Coefficients from univariate Cox regression were z-scored. Established prognostic signatures were compiled by CIDE. **g,** Bar plots showing the relative predictive power of gene signatures for PFS (top) and OS (bottom) in ipilimumab-pretreated, anti-PD-1-treated Liu2019 patients. **h,** Kaplan–Meier curves showing PFS (top) and OS (bottom) in ipilimumab-pretreated, anti-PD-1-treated Riaz2017 patients stratified by GRN scores. P-value, two-sided Wald test for the signature term from univariate Cox regression. **i,** Bar plots showing the relative predictive power of gene signatures for PFS (top) and OS (bottom) in ipilimumab-pretreated, anti-PD-1-treated Riaz2017^91^ patients. **j,** Error-bar plots showing hazard ratios and 95% confidence intervals (CIs) for the GRN signature in predicting PFS (top) and OS (bottom) in integrated ipilimumab-pretreated, anti-PD-1-treated Campbell2023^90^ cohorts. P-value, two-sided Wald test for the signature term from univariate Cox regression. **k,** Forest plots showing hazard ratios and 95% CIs for multiple terms in predicting PFS (top) and OS (bottom) in integrated ipilimumab-pretreated, anti-PD-1-treated Campbell2023 cohorts. P-values, two-sided Wald tests from Cox regression. Box plots show median and interquartile range, with whiskers extending to 1.5× IQR.

Together, these analyses demonstrate that key melanoma gene regulatory modules are stably maintained in vitro, faithfully reflect tumor-TME co-adaptation in vivo, and link to immunotherapy responsiveness. Integrating single-cell and bulk transcriptomic data across model and patient contexts highlights how multi-omic dissection of cancer gene regulation can advance precision oncology.

## Discussion

The complexity of cancers has long hindered a unified understanding of their underlying molecular features and the development of targeted therapeutics. Here, we leveraged a comprehensive cancer cell line resource to dissect gene regulatory principles and convergent molecular features of cancer. Using our custom EasySci single-cell platform with highly parallel inter-sample processing capacity^13,95^, we performed single-cell transcriptomic and epigenetic profiling of 60 cell lines spanning 16 tissue origins and 24 cancer types. Compared with the previous study that profiled single-cell transcriptomes and chromatin accessibility across cancer cell lines^10^, we expanded RNA profiling by more than 10-fold and obtained over 4-fold more single-cell ATAC profiles, covering more tissue origins and cancer types and enabling a more unbiased characterization of intra- and inter-lineage variation.

Beyond prior single-modal examinations of transcriptomic phenotypes in cancer cells^12,96^, our cross-modal dataset enabled integrative analyses linking chromatin landscapes with transcriptomes to identify gene regulatory phenotypes driven by differential activation of TF-regulated gene expressions^97^. The observation that these GRN-based cancer cell line clusters either reflect tissue origin and cancer type or contain highly heterogeneous cell lines is consistent with previous pan-cancer integrative clustering on bulk patient samples across copy number, DNA methylation, mRNA, and miRNA profiles^98^, in which both single cancer-type-dominant and mixed clusters were identified. This reveals the coexistence of tissue-specific molecular programs reflective of developmental origins and convergent features shared across lineages. The diversity we observed in cell lines further supports the maintenance of key transcriptional phenotypes of cancer under in vitro conditions, highlighting their utility for drug-response measurement and mechanistic dissection^99^.

Regarding phenotypic convergence across cancers, extensive characterization of cross-lineage cancer cell line clusters supports their reflection of EMT^25,100^, reinforcing the existence of a universal transcriptional dedifferentiation program across epithelial cancers, with EMT representing the dominant cross-lineage phenotypic spectrum. Although the universality of EMT across cancer types has been widely debated^34,101,102^, our de novo identification of this phenotypic spectrum at the pan-cancer level provides strong supporting evidence. Moreover, while a few TFs such as ZEB1 have long been recognized as essential EMT drivers^25^, recent evidence suggests a more complex mode of action involving cooperative regulation by multiple TFs, including AP-1, TEADs, and ZEB1^36^. The incomplete phenotypic reversal observed upon inhibition of single TFs further supports the involvement of additional factors^103,104^. Beyond ZEB1, FOSL1, and FOSL2 identified here, other strong EMT-associated TFs nominated from our study may expand this repertoire, offering potential avenues for combinatorial inhibition. Conversely, the reidentification of epithelial-state-associated transcription factors, including GRHL2, CEBPB, and FOXC1, further supports the convergence of EMT-associated regulatory programs across cancer types. While these factors have been implicated in EMT regulation in specific cancer contexts^37–39^, our pan-cancer single-cell analysis places them within a broader cross-lineage framework, highlighting recurrent epithelial-associated regulatory features shared across diverse cancer cell states. These analyses provide additional mechanistic insight into the regulatory dynamics underlying cancer-associated EMT.

Previous studies have widely identified TF amplification as a key event in carcinogenesis^105^; for example, MYC focal amplification through ecDNA formation enhances cellular fitness under microenvironmental stress^106^. Recent work exemplified how chromosomal amplification can alter cancer cell states through TF overactivation, as shown in a case study of SOX4 in glioblastoma subclones^9^. Here, we systematically identified preferential amplification of TFs, rather than their target genes, as a common phenomenon across cancers. Our association and validation analyses further nominated CEBPB as a TF that enhances the epithelial state through copy number amplification, thereby extending this TF amplification-driven regulatory mechanism to more benign cell states. Moreover, our analyses nominated another set of genes whose amplification may constrain EMT through transcription-independent mechanisms. Similarly, a previous study identified tumor-suppressive amplification of chromosome 21 linked to hyperactivity of calcineurin suppressors^107^. Such mechanistic diversity underscores the need for caution in therapeutic design despite the convergence of cancer cell states^108^.

Given the abundance of melanoma cell lines in our cohort, we performed extensive comparative analyses to identify gene regulatory features specific to AM relative to the more common cutaneous subtype (CM). Only limited comparisons have been made between these subtypes, primarily focusing on genomic alterations^64^ and their therapeutic outcomes^109^. Complementarily, we explored their differences in gene regulation. By deconvolving gene programs and constructing gene regulatory networks for each subtype, we found that JAK-STAT signaling activity and inflammatory transcriptional responses represent the primary distinctions. Notably, many TFs we identified as melanoma subtype-specific are also key regulators of melanoma phenotypic switching, including SOX10 and MITF in melanocytic differentiation and FOSL1 and JUN in the development of the undifferentiated state^24,110^. As our orthogonal gene program deconvolution presents these subtype-independent phenotypic states in parallel, the enrichment of these TFs within subtype-specific regulatory architectures strongly suggests their multifaceted roles in tuning coexisting phenotypes in melanoma.

Lastly, by extrapolating gene regulatory features derived from cancer cell lines to their corresponding TME characteristics, we found that the AM gene program is associated with an immune-suppressive TME composition, consistent with prior spatial transcriptomic studies of AM showing a globally low level of immune infiltration^80^. Through integrated analysis of patient-derived TME cell-type-specific transcriptomes and melanoma cell lines, we identified TGF-β1 as a potential key reprogramming factor shaping the AM TME, whereas CM was characterized by active anti-tumor immunity and a potentially heightened IFN response. As previously reported, TGF-β signaling robustly attenuates tumor response to PD-L1 blockade^111^, suggesting its potential as a therapeutic target in the context of AM. Furthermore, our subtype-specific gene regulatory network successfully classified patients by immunotherapy response, consistent with observations from large clinical cohorts showing that AM exhibits lower response rates and poorer prognosis than CM^109,112^.

Although we profiled 60 cell lines spanning 20 cancer types, our panel does not encompass the full diversity of human cancers. Several clinically important malignancies, such as pancreatic, gastric, and bladder carcinomas, were not represented, and therefore the extent to which the regulatory programs identified here generalize to these contexts remains to be determined. Future studies incorporating broader cancer type coverage, additional genetic backgrounds, and primary tumor or patient-derived models will be important for defining which regulatory programs are broadly conserved versus lineage- or context-specific.

A second limitation is that our study profiles cancer cell lines cultured in isolation, which do not fully recapitulate the complex tumor microenvironment, including immune, stromal, endothelial, and fibroblast components, as well as the extracellular matrix. Therefore, our dataset is best interpreted as a resource for dissecting cancer cell-intrinsic regulatory programs. Nevertheless, cell lines provide a tractable and controlled system that minimizes confounding from variable tumor purity, inter-patient heterogeneity, and tissue composition, factors that can complicate analyses of primary tumors, especially for rare cancer subtypes with limited available specimens. The retention of key phenotypic states and regulatory programs in vitro supports the utility of these models for studying intrinsic cancer cell states. Although our analyses suggest that the AM- and CM-associated programs primarily reflect melanoma-cell-enriched transcriptional states, variation in tumor purity and TME composition may still contribute to program scoring in bulk tumors.

In addition, ligand-receptor inference should be interpreted as an exploratory analysis for hypothesis nomination. Our patient-level tumor-microenvironment inferences also relied primarily on a single acral melanoma single-cell cohort with limited clinical annotation, and larger, more completely annotated cohorts will be needed to assess how ethnicity, treatment history, and other clinical variables shape TME composition and cell states. Future integration with subtype-resolved single-cell, spatial transcriptomic, or imaging-based datasets will help further distinguish malignant-cell-intrinsic programs from microenvironment-associated components and strengthen mechanistic understanding of how specific TME phenotypes are established and maintained.

Overall, this single-cell multi-omics resource and our analyses comprehensively delineate pan-cancer gene regulatory principles and reveal diverse modes of altering cancer cell states. By uncovering the TME-coadapted gene regulatory landscape of melanoma cell lines in vitro, our findings highlight the fidelity of in vitro cancer models in recapitulating clinically relevant regulatory programs and point to the potential of transforming molecular multi-omics insights into clinical interpretation and translational application.

## Methods

### Cell culture

For the acral melanoma cell lines: YUHIMO, YUSEEP, and YUSUSA were obtained from Yale University. M040204, M040416, M141207, and M160113 were obtained from the University of Zurich. WM3211, WM4325, and WM4324 were obtained from the Wistar Institute. MB4667 was obtained from the University of Colorado. The remaining cell lines (HCC1954-LCC1, HCC1954-LCC2, MDA-231-BrM2-831, MDA-231-LM2-4175, MDA-231-AdM-1834, MDA-231-BoM-1833, MDA-231-TGL, SK-BR-03, H2030-BrM3, H2030-TGL, H2087-TGL, H2087-LCC1, H2087-LCC2, PC9-BrM3, PC9-TGL, Calu-1, SK-LC-17, SH-SY5Y, SK-N-AS, 786-M1A, 786-M2B, 768-O-TGL, CAPAN02, LNAR, LNCAP-EGFP, LNCAP-EGFP-c.2-F876L, HT-29, SK-CO-01, SK-OV-03, SK-UT-01, SK-HEP-01, MSK921, OS252, SK-ES-01, and SK-NEP-01 cell lines) were obtained from the Antibody and Bioresource Core Facility at Memorial Sloan Kettering Cancer Center. IMR90, MeWo, RPMI7951, Sk-Mel-24, Sk-Mel-3, DB, GA-10-Clone-4, MOLT-4, HL-60, U2-OS, and MP41 cell lines were obtained from the American Type Culture Collection. All cell lines were maintained at 37 °C with 5% CO2.

### EasySci RNA Library Preparation

Libraries were prepared following the published EasySci-RNA protocol^13^. Nuclei were isolated from cultured cell lines and distributed at up to 20,000 nuclei per well across two 96-well plates in Nuclear Suspension Buffer. Reverse transcription used dual priming (50 µM short-dT and 50 µM randomN) with Maxima H- Minus Reverse Transcriptase and a staged temperature ramp (55 °C denaturation followed by 4–55 °C gradient). Post-RT, nuclei were pooled, washed, and redistributed for plate-specific barcoding by DNA ligation with pre-annealed ligation adapters. Second-strand synthesis (NEB Ultra II Non-Directional) was performed, followed by 0.8× AMPure cleanup. Libraries were then tagmented with loaded Tn5, quenched with SDS/BSA, and indexed by addition of universal P5 and indexed P7 primers, followed by amplification with NEBNext High-Fidelity 2× Master Mix (12–15 cycles determined by qPCR when applicable). Final libraries were purified, quantified by Qubit, size-checked on 2% agarose, and sequenced on an Illumina NextSeq 1000 (paired-end 100 bp; Index 1/2: 10 bp each).

### EasySci ATAC Library Preparation

Libraries were prepared following the published EasySci-ATAC protocol^13^. In brief, nuclei were isolated from cultured cell lines and approximately 5,000 nuclei per reaction were subjected to indexed Tn5 transposition in 20 mM Tris-HCl, 20 mM MgCl₂, and 20% DMF for 30 min at 37 °C, followed by quenching with 1× EDTA stop buffer. Tagmented nuclei were pooled, redistributed across 96-well plates, and barcoded via plate-specific P5 ligation. After pooling and cleanup, libraries were amplified with indexed P5/P7 primers using NEBNext High-Fidelity 2× PCR Master Mix. Final libraries were purified with AMPure XP beads, quality-checked by agarose electrophoresis and Qubit quantification, and sequenced on an Illumina NextSeq 1000 (100-cycle kit; Read 1: 58 cycles, Read 2: 60 cycles, Index 1/2: 10 cycles each).

### Sequencing Data Processing

We used custom computational pipelines for processing EasySci-RNA and -ATAC data as previously described in^13^. In brief, for the RNA modality, UMI and cell barcode sequences of each read pair were extracted from specific locations within the reads. Adapter trimming on Read 2 was performed using Trim Galore with default settings to remove poly(A) sequences and low-quality bases from the 3′ end. The paired-end sequences were then aligned to the genome with STAR, and PCR duplicates were removed using both UMI and genomic coordinate information. Reads were subsequently split into single-cell SAM files, and gene expression counts were generated.

For the ATAC modality, N5, N7 Tn5 barcodes and cell barcodes were extracted, and paired-end reads were aligned with STAR after adapter trimming with Trim Galore. Only uniquely aligned read pairs were retained, and PCR duplicates were removed using Picard at the PCR-well level. BED files containing mapped fragment start and end sites, strand information, and cell barcodes were generated for downstream processing.

Scanpy (v1.11.0)^113^ was used for single-cell RNA-seq analysis. Scrublet (v0.2.3)^114^ was applied for doublet identification. To visualize RNA profiles of single cells, gene counts were normalized and log1p transformed, followed by selection of the top 5,000 highly variable genes. Regression on total cell counts and subsequent scaling of HVG expression across cells were performed. The top 30 principal components (PCs) were then used to construct the nearest neighbor graph and the global UMAP.

SnapATAC2 (v2.8)^115^ was used for single-cell ATAC-seq fragment demultiplexing and downstream analyses. PCR-well level BED files were indexed and used as inputs. Only cells with at least 1,000 fragments and a TSS enrichment (TSSE) score ≥ 3 were retained. Peak calling was performed at the cell line level using the macs3 function, and peaks overlapping with the ENCODE human genome blacklist^116^ were removed. Significant peaks (q < 1e−3) were iteratively merged. The single-cell ATAC peak count matrix was constructed using the make_peak_matrix function, and the global UMAP was computed using the top 30 spectral embeddings derived from the 300,000 most variable peaks.

### Single-cell RNA-ATAC integration

Single-cell RNA-ATAC integration was performed using GLUE (v0.3.2)^117^. Because our pan-cancer cell line cohort included multiple cell lines with high transcriptomic similarity, integration was carried out at the individual cell line level. In brief, for each cell line, a preprocessed AnnData object containing a raw single-cell gene expression count matrix with the top 3,000 highly variable genes and PCA embedding, and an ATAC AnnData object containing the spectral embedding from the top 300,000 peaks were generated using Scanpy and SnapATAC2, respectively, and provided as input to GLUE. A guidance graph was constructed based on the genomic proximity of ATAC peaks to annotated genes using the reference genome annotation. To improve integration quality, Leiden clustering and UMAP visualization were performed. Clusters consisting predominantly of cells with low RNA UMI counts, low ATAC fragment counts, or low TSSE scores were removed, and a second round of integration was performed using the refined cell sets. Integration quality was evaluated both manually and computationally. Manual assessment was conducted by visualizing cell cycle scores in the GLUE-derived co-embedding UMAP, while integration consistency scores from GLUE were computed in parallel to provide a quantitative measure of integration accuracy.

### Multi-modal meta-cell identification

Multi-modal meta-cell identification was performed using SEACells (v0.3.2)^118^. For each cell line, the final GLUE co-embedding was used to construct kernels, and meta-cells containing at least 5 cells from both modalities, with each modality contributing 20–80% of the total cell number, were retained for downstream analyses. The expected number of meta-cells per cell line was estimated as (total cell number)/75. For each meta-cell, raw gene expression counts and ATAC peak counts from all assigned cells were aggregated. For cell lines with poor integration quality, specifically, those with co-embeddings that failed to capture clear cell cycle progression (often associated with low overall UMI counts), or with severe imbalance between RNA and ATAC cell numbers (log2(RNA cell number / ATAC cell number) > 3 or < –3), all cells were collapsed into a single meta-cell.

### Pan-cancer gene regulatory network construction and regulon-based analyses

Pan-cancer–level TF–cis-regulatory element–target gene triplets were identified using SCENIC+ (v1.0a2)^119^. In brief, meta-cell RNA profiles were preprocessed with Scanpy, and the raw count matrix was stored in the .raw slot. Meta-cell ATAC profiles were preprocessed using pyCisTopic^120^, and the top 35 topics were selected based on the diagnostic metrics implemented in the package. Candidate peaks were defined by combining (i) the top 3,000 peaks with the maximum probability of belonging to each topic, (ii) peaks identified within each topic using Otsu thresholding, and (iii) highly variable peaks that were significant markers of each cell line. A custom cisTarget motif library was then built using candidate peaks with ±1 kb extension as background padding. Linkage identification and AUCell scoring at the meta-cell level were performed using the SCENIC+ Snakemake pipeline.

Given the more established role of TFs in transcriptional activation and the greater number of activation regulons, we retained only multimodal regulons showing positive associations between TFs and target genes, as well as between ATAC peak accessibility and target genes (+/+). For each cell line, the mean AUCell score of each regulon was calculated, and hierarchical clustering was performed to identify GRN clusters across cell lines.

To assess cell-line–specific regulon activation, Wilcoxon rank-sum tests were performed on regulon scores (computed with the score_genes function in Scanpy) between meta-cells from the focal cell line and the remaining meta-cells. For cell lines that had been pseudobulked into a single sample for GRN construction, RNA-only meta-cells were generated by randomly sampling 50 cells per meta-cell. Regulons were considered active in a given cell line if they passed an FDR threshold of 0.01 and exhibited a mean regulon score at least 0.1 greater than background.

### Differential gene expression and differential peak accessibility analyses and functional enrichments

The pan-cancer cell line multi-modal marker examination was conducted using the rank_genes_groups function in Scanpy. Only genes expressed in at least 200 cells were retained for the analysis. Marker examinations on ATAC were performed similarly using the normalized, log1p-transformed gene activity, which was obtained by summarizing the ATAC signal coverage across the gene body regions. The top significant (FDR < 1e-3) cell line markers with the highest log2 fold changes in both modalities were further intersected.

To further include the single-cell data sparsity into consideration and minimize the false positive discovery^121^, DESeq2^58^ was used to examine GRN-cluster-specific differentially-expressed genes at the meta-cell level. As there is no single universal differentially accessible peak identification method that consistently outperforms the rest of them, the marker_regions function in snapATAC2 was used to identify GRN-cluster-specific differentially accessible peaks. Only peaks with counts in at least 2.5% cells and showing absolute log fold change >= 0.25 were considered. Functional enrichment on DEGs was conducted using Enrichr^122^; GREAT was used for functional enrichments on genomic regions^123^.

### EMT trajectory inference and trajectory-associated regulon analysis

A curated gene list of conserved upregulated and downregulated genes during EMT, identified from cytokine-treated cell line panels^34^, was used to order cells in low-dimensional space. PCA on these genes across all RNA meta-cells was performed, and the root of the transition trajectory was determined by scoring these genes. Slingshot (v2.16.0)^40^ was then applied to infer pseudotime along the trajectory.

To identify pan-cancer regulons associated with EMT, generalized additive models (GAMs) were fitted using the gam() function from the mgcv R package, modeling regulon gene program scores as a function of meta-cell pseudotime. Each GAM was compared to a null model using the anova() function, and regulons with significantly improved fits were considered pseudotime-dependent. To evaluate directionality, pseudotime was divided into 30 bins, and the mean regulon scores were correlated with mean pseudotime across bins. Background distributions were generated by shuffling pseudotime across meta-cells. Regulons with Pearson correlation coefficients > 0.4 or < -0.4 and FDR < 0.01 were designated as positively or negatively associated, respectively. As an additional filter, the expression of the TF itself was modeled against pseudotime using GAM; only regulons whose TFs showed significant pseudotime dependence and had correlations > 0.4 between TF expression and regulon score were retained as final regulators.

ATAC motif deviations of TFs were computed using chromVAR (v1.30.1)^124^. Motif matches within ATAC peaks were identified using the matchMotifs() function, and motif deviations were computed relative to GC-matched background peaks using the computeDeviations() function. To ensure comprehensive coverage, position weight matrices (PWMs) of motifs included in the SCENIC+ GRN construction but absent from the default JASPAR database were manually merged into a custom supplementary motif database for chromVAR.

### Copy number variation inference and association analyses

InferCNV (v1.24.0)^125^ was used to call CNVs at the meta-cell level based on RNA raw counts. We adopted a reference-free mode to avoid background overcorrection. Gene coordinate files were generated from the reference genome annotation used for read mapping, and RNA meta-cells were grouped by cell line identity. Genes were ordered by genomic position and normalized expression values were smoothed across neighboring genes within each chromosome to identify broad regional expression shifts relative to the cohort-wide pseudo-reference baseline. Then the hidden Markov model (HMM) of inferCNV, i6 HMM model, produced six CNV states: 0 = complete loss, 0.5 = single-copy loss, 1 = neutral, 1.5 = single-copy gain, 2 = two-copy gain, and 3 = more than two-copy gain. Bayesian post-filtering of HMM-defined CNV regions was performed using a default posterior normal-probability cutoff of 0.5, and genomic regions with CNVs supported by posterior probabilities > 0.5 were retained. Retained CNV events were further evaluated based on regional consistency and orthogonal ATAC/WGS/WES support.

To identify associations between EMT status and gene-level CNVs across pan-cancer cell lines, Pearson correlation coefficients were computed between pseudotime-inferred EMT status and gene copy number states derived from the HMM model. Multiple testing correction was performed across all genes using FDR.

To further validate “hotspot” regions associated with EMT, ATAC-seq coverage across gene bodies was extracted, and correlations with EMT status were computed in the same manner. Only genes that showed consistent and significant correlation trends at both the RNA-inferred CNV and ATAC coverage levels were considered.

### Validation analysis on CEBPB by in-silico knockout and ChIP-seq

SCENIC+^119^ was used to simulate the perturbation effects of CEBPB. In brief, target genes of each regulon together with the RNA meta-cell count matrix were input into the train_gene_expression_models() function to construct random forest models, modeling the expression of target genes as a function of upstream regulators. CEBPB expression was then set to zero, and 10 iterations of perturbation simulation were performed using the simulate_perturbation() function. The resulting simulated transcriptomic states with CEBPB expression eliminated were projected onto the EMT PCA embedding of meta-cells using the plot_perturbation_effect_in_embedding() function.

To map additional cell lines (MCF7, HeLa, A549, HepG2, HCT116) onto the EMT trajectory, we reprocessed single-cell RNA-seq data for these lines^10^, which were selected because ENCODE^53^ provides CEBPB ChIP-seq profiles for them. To minimize potential biases from data sparsity differences, single cells from these datasets were randomly sampled and aggregated into meta-cells of 20 cells each, given their deeper sequencing depth. After normalization, meta-cell count matrices from the internal and external datasets were combined, retaining only genes from the conserved EMT gene sets described above. Genes were then scaled across all meta-cells. The get.knnx() function from the FNN R package was used to identify the two nearest neighbors of each external meta-cell within the scaled internal meta-cell matrix. The mean EMT pseudotime of these two neighbors was assigned to the external meta-cell, and the median transferred EMT pseudotime was used to represent the EMT status of each external cell line. Finally, CEBPB-to-background p-value signal tracks from ENCODE ChIP-seq experiments were visualized using the IGV tool (v2.19.6)^126^.

### Gene program deconvolution and signaling activity scoring on skin cancer cell lines

The oNMF module of D-SPIN (v1.4.0)^57^ was used to deconvolve orthogonal gene programs from single-cell RNA-seq profiles of skin cancer cell lines. Manual curation of gene program members was performed, and the number of programs chosen was determined based on those yielding the most biologically interpretable results. To further refine core gene programs, differential gene expression analyses on meta-cells were conducted in parallel using DESeq2. For programs 4 and 11 (AM/CM universal programs), member genes were intersected with significant DEGs between AM and CM cell lines (FDR < 0.01). For other AM/CM cell line–specific programs, member genes were intersected with significant DEGs comparing the given cell line to all other cell lines (FDR < 0.01). Meta-cell program scores were computed using the score_genes function in Scanpy.

Signaling activities of meta-cells were estimated using PROGENy (v1.17.3)^127,128^, which infers pathway activity by weighting predefined “footprint” genes. Hormone-associated pathways (estrogen and androgen) were excluded from analysis.

### Examination of AM/CM-specific transcriptional regulation

For finer-grained skin cancer subtype–specific GRN construction, we constrained the set of genes and ATAC peaks examined in FigR (v0.1.0)^129^. Only genes from DE-refined program 4 and program 11, and peaks showing significantly different accessibility between AM and CM cell lines (identified using SnapATAC2; only peaks detected in ≥1% of cells and with absolute log fold change ≥ 0.25 were considered) were used for peak–gene association. For each candidate gene, 100 permutations were performed to compare the observed peak–gene correlation with correlations generated from GC-matched background peaks. Genes with at least four associated peaks were further analyzed for TF regulation by testing (i) TF motif over-representation in the associated peaks and (ii) correlation of candidate TF expression with accessibility of these peaks. Only positive TF regulators whose expression significantly correlated with peak accessibility (p < 0.1) and whose motifs were significantly enriched in the same peaks (p < 0.1) were retained as master regulators of the peaks and their downstream target genes.

As inputs included only AM-specific genes (DE-refined program 11) and CM-specific genes (DE-refined program 4), we further quantified TF specificity. For each TF, the proportion of AM- or CM-specific target genes among all AM- or CM-specific genes targeted by all TFs was calculated. Proportion tests with FDR correction across all TFs were then conducted to identify AM- and CM-specific TFs exerting significantly stronger regulation on larger sets of target genes in a given skin cancer subtype.

### Bulk and single-cell RNA-seq TME analyses

Bulk RNA-seq count files from the melanoma cohort of TCGA^130^ were downloaded by selecting Program: TCGA, Project: TCGA-SKCM, and Primary Site: skin. Bulk RNA-seq–based tumor microenvironment (TME) immune cell type deconvolution results were obtained from TIMER2.0^131^. Patients’ bulk RNA-seq profiles were scored against DE-refined skin cancer gene programs using the AddModuleScore() function in Seurat^132^. Pearson correlations were computed between program scores and TME cell type abundances across patients, followed by FDR correction of correlation p-values. Predicted neoantigen counts derived from TCGA patient SNV profiles were obtained from^78^. Patients were grouped and Wilcoxon tests were applied to compare metrics across groups.

Single-cell RNA-seq datasets profiling the TME of both AM and CM patients were obtained from^60^. Patient-level AnnData objects were concatenated, and standard single-cell processing was performed, including normalization, log1p transformation, highly variable gene selection, regression on gene number, PCA, nearest-neighbor graph construction, UMAP embedding, and Leiden clustering. Melanoma cell clusters characterized by high MITF expression, as well as patient-specific clustering patterns, were removed. TME cell types in MITF- or PTPRC+ clusters with contributions from multiple patients were retained for an additional round of processing and doublet removal using doubletdetection^133^. Clusters with doublet proportions >30% were excluded. Final cluster annotation was performed using a combination of representative marker gene expression and CellTypist^92^, an automated cell type annotation tool. Key immune cell phenotypic signatures were scored using starCAT (v1.0.9)^82^, and comparisons between AM and CM conditions were conducted using Wilcoxon tests followed by FDR correction. Differential expression results for key functional markers and ligand/receptor genes were extracted from global differential expression analyses of each TME cell type between AM and CM conditions. Ligand/receptor genes were selected from the KEGG database^134^ under the term "CYTOKINE CYTOKINE RECEPTOR INTERACTION".

Patient immunotherapy response and survival data were obtained from the Cancer Immunology Data Engine (CIDE)^87^. AM target genes of AM-specific TFs were used as negative biomarkers, while CM target genes of CM-specific TFs were used as positive biomarkers. Biomarker scores for patients were computed using pretreatment transcriptomic data from the database, and these scores were used for patient stratification.

## Acknowledgments

We thank the members of the Cao Lab at Rockefeller University for their generous feedback. We also thank the members of the Rockefeller University High-Performance Computing Core, Genomics Resource Center, and Bioinformatics Resource Center for their technical support. The schematic in Fig. 1 was created with BioRender.com. We are grateful to Dr. Kasey Couts (University of Colorado), Dr. Meenhard Herlyn (Wistar Institute), Dr. Mitch Levesque (University of Zurich), and Dr. Ruth Halaban (Yale University) for kindly providing Acral Melanoma cell lines.

## Author contributions

J.C. and W.Z. conceived the study. A.U. and J.K. performed cell culture and EasySci-ATAC library preparation. C.S. performed EasySci-RNA library preparation. A.U. conducted sequencing read preprocessing and organization. Z.X. performed computational analyses with input from all co-authors. Z.X., J.C., W.Z., and A.U. wrote the manuscript with input from all co-authors.

## Data availability

Raw FASTQ files generated in this study have been deposited under NCBI BioProject accession PRJNA1354039. Processed gene count matrices, ATAC peak count matrices, and cell metadata are available in the NCBI Gene Expression Omnibus (GEO) under accession number GSE311521. Public reference lung and breast tissue single-cell RNA-seq atlases data were retrieved via CZ Cell x Gene portal (https://cellxgene.cziscience.com/). Public CNV and drug sensitivity data for cell lines were obtained from the Cancer Cell Line Encyclopedia (CCLE, https://sites.broadinstitute.org/ccle/). Public ChIP-seq data were obtained from ENCODE (https://www.encodeproject.org/). TF-ome CRISPRa Perturb-seq on RPE-1 cell line was obtained from Zenodo (https://zenodo.org/records/15213619); Genome-wide CRISPRi FiCS Perturb-seq on HEK293T cell line was obtained from Figshare (https://doi.org/10.25452/figshare.plus.29190726.v2). Public single-cell RNA-seq on patients’ TME and bulk RNA-seq on patient cancer samples were obtained from GSE215121 of GEO. Public immunotherapy patient cohort data were collected by CIDE (https://cide.ccr.cancer.gov/).

## Code availability

Analysis scripts used in this study are available at GitHub (https://github.com/JunyueCaoLab/Cancer_atlas).

